# The microbiomes of seven lichen genera reveal host specificity, a reduced core community and potential as source of antimicrobials

**DOI:** 10.1101/789032

**Authors:** Maria A Sierra, David C Danko, Tito A Sandoval, Gleb Pishchany, Bibiana Moncada, Roberto Kolter, Christopher E. Mason, Maria Mercedes Zambrano

**Author notes:** Corresponding author: Maria M. Zambrano, Corporación Corpogen, Carrera 4 # 20-41, Bogotá DC, 110311, Colombia.

## Abstract

The High Andean Paramo ecosystem is a unique neotropical mountain biome considered a diversity and evolutionary hotspot. Lichens, which are complex symbiotic structures that contain diverse commensal microbial communities, are prevalent in Paramos. There they play vital roles in soil formation and mineral fixation. In this study we analyzed the microbiomes of seven lichen genera in two Colombian Paramos using 16S rRNA gene analyses and provide the first description of the bacterial communities associated with *Cora* and *Hypotrachyna* lichens. Paramo lichen microbiomes were diverse, and in some cases were distinguished based on the identity of the lichen host. The majority of the lichen-associated microorganisms were not present in all lichens sampled and could be considered transient or specialists. We also uncovered sixteen shared taxa that suggest a core lichen microbiome among this diverse group of lichens, broadening our concept of these symbiotic structures. Additionally, we identified strains producing compounds active against clinically relevant pathogens. These results indicate that lichen microbiomes from the Paramo ecosystem are diverse and host-specific but share a taxonomic core and can be a source of new bacterial taxa and antimicrobials.

## Introduction

Symbiotic relationships between eukaryotes and microorganisms are ubiquitous (1), and often essential for the function and survival of the host, fulfilling roles that range from stress tolerance (2) and nutrient supply (3, 4) to defense against pathogens (5, 6). The composition of the microbial community associated with a particular host is defined by factors such as temperature and pH (7), host genotype, nutrients (8) and microbe-microbe interactions (9). Recent evidence indicates that hosts of the same species (8), as well as evolutionarily related hosts (10), harbor similar microbial communities. Core microbiomes, which are microbes consistently associated with a given host or found in a large fraction of samples from a particular environment(11), have been identified and catalogued in sponges (7), corals (12), insects (13), plant roots (14) and mammals (10, 15).

Lichens represent some of the oldest and most diverse symbioses on Earth (16). Lichens consist of a photobiont (cyanobacterium/alga) and a mycobiont (fungus), which together form a unique structure called the thallus (17). Lichens play a vital role in ecosystems as they are essential in soil formation, naked soil colonization, and nutrient uptake and release for plants (18, 19). Lichens can colonize a wide range of substrates, from natural surfaces to man-made materials such as plastic, rubber, metals and glass (20). They can also tolerate extreme environmental conditions and offer a niche for diverse microorganisms (21, 22). The diversity of these lichen-associated microbial communities is not yet well characterized, and has only recently been investigated using high-throughput techniques (23, 24). These studies indicate substantial microbial and functional diversity (24-26) that has been suggested to help protect the thalli against pathogens through the production of antimicrobials (27, 28). The process of community establishment within lichens is poorly understood and has been proposed to be driven either by the photobiont (29) or by geography and habitat (30).

Recent studies have described the microbial communities associated with lichens using culture-independent strategies (24, 25). However, comparisons between studies are hindered by differences in sampling methods, data analyses, and poor or complete lack of lichen description. Given the complexity of the lichen symbiotic structures, with recent evidence indicating that some lichens may be composed of multiple bacteria and more than one fungus (26), it is important to study lichen microbiomes in order to understand their ecological role in the symbiosis.

A large and unexplored diversity of lichens is located in the Paramo ecosystem (31), a unique biodiversity hotspot that consist of high-elevation regions distinguished by extreme daily temperature variations, nutrient-poor and acidic soils, heavy rains and high UV radiation (32). The Paramo, as many of Colombia’s Andean ecosystems, has been understudied for decades due to armed conflict (33). Here we characterize and compare the microbial communities in seven lichen genera from two Colombian Paramos. Using amplicon sequencing of the 16S rRNA gene, we describe these microbiomes and identify members common to all seven genera, expanding our understanding of the complexity underlying these symbiotic structures. In addition, we isolated bacteria producing antibacterial and antifungal compounds and propose that Paramo lichen microbial communities should be further studied as a valuable source of antimicrobials. The diversity and prevalence of lichen-associated microbial communities underscore the need the further understand their ecological roles in lichen function.

## Results

### Microbial diversity varies depending on the lichen host

To study the structure of bacterial communities in lichens, we collected samples from different lichen genera at Paramo ecosystems within two national parks in Colombia. The 57 lichen samples were classified into eleven genera (Table S1), but only seven genera (*Cora, Hypotrachyna, Usnea, Cladonia, Peltigera, Stereocaulon* and *Sticta*), which corresponded to 47 individual samples, were found in both locations. Samples with three or more biological replicates from the seven genera were used for microbial community analyses. DNA was isolated from individual lichen samples to identify microbial community profiles by 16S rRNA sequencing, which resulted in a total of 3,412,279 reads (mean per sample: 72,601).

A total of 20,174 operational taxonomic units (OTUs) were identified across all samples, which ranged from 100 to 1,955 OTUs per sample. Rarefaction curves indicated that this richness was adequately sampled as many samples reached saturation (Figure S1). Simpson and Shannon diversity indices were calculated after randomly subsampling to the lowest number of reads (20,623), showing a broad distribution among samples. Diversity was significantly different between *Usnea* and *Hypotrachyna* lichens (Figure 1a and Table S2, using ANOVA (p=0.037 and p=0.026 for Simpson and Shannon indices, respectively), and between lichens *Usnea* and *Sticta,* which had the smallest and largest number of OTUs, respectively (ANOVA test p=0.010).

**Figure 1.**
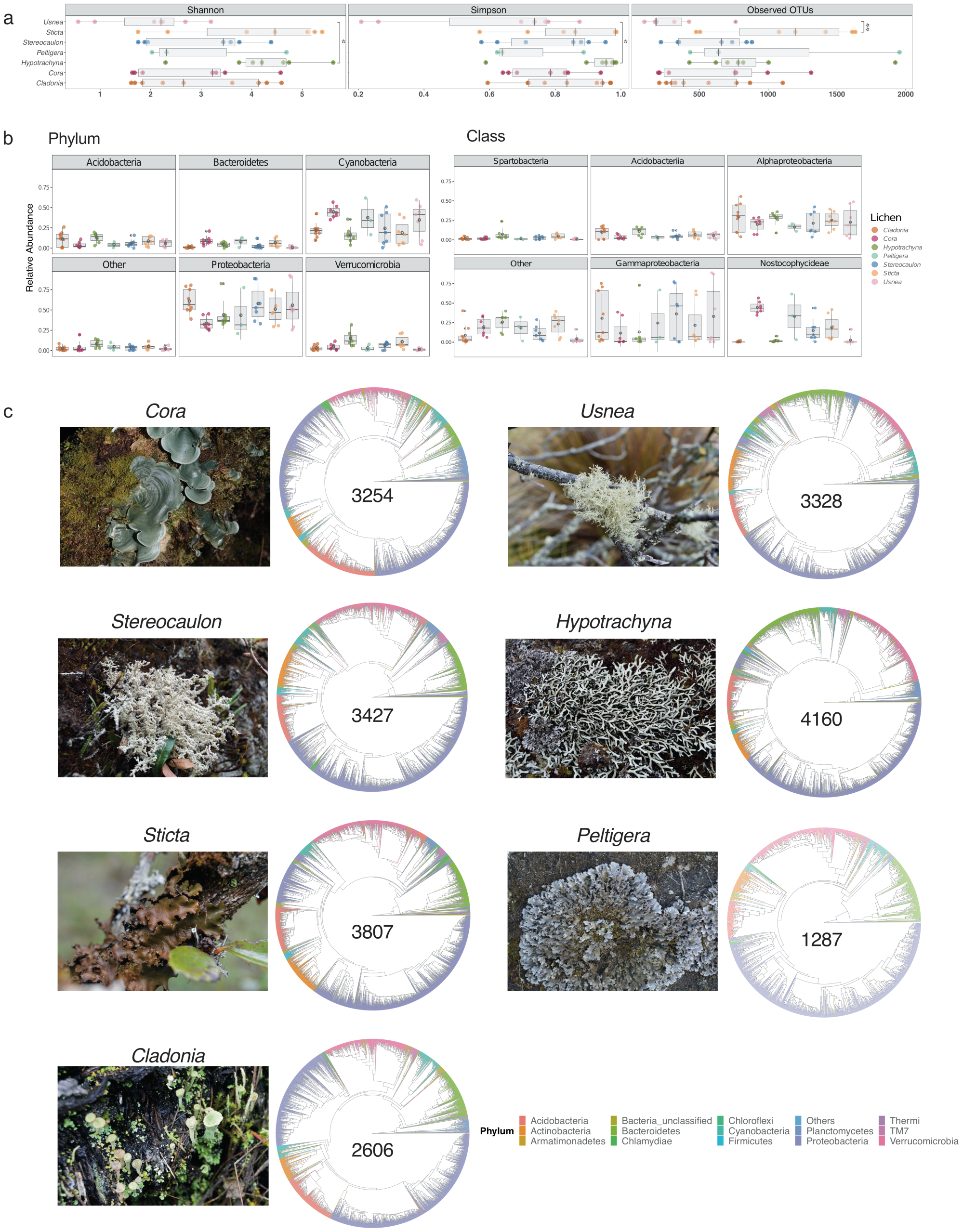
Microbial diversity varies across seven lichen genera. **a**. Diversity and richness values for samples per lichen genus, measured by the Shannon and Simpson indices and the number of Observed OTUs, respectively. Multiple comparisons of richness and diversity measures were performed by one-way ANOVA with P values of <0.05 considered to be statistically significant. Asterisks indicate significant differences between *Usnea* and *Hypotrachyna* for Simpson (p=0.037) and Shannon (p=0.026) indices, and between *Usnea* and *Sticta* (p=0.01) for Observed of OTUs. **b**. Relative abundance of the five most abundant phyla and classes. Data are presented as mean ± standard deviation (SD). **c**. Taxonomic diversity of lichens with the total number of OTUs for a given genus (shown in the center of cladogram).

Taxonomic assignment of OTUs showed that lichen microbiomes were predominantly composed by members of the phyla Acidobacteria, Actinobacteria, Bacteroidetes, Cyanobacteria, Proteobacteria and Verrucomicrobia (Figure S2). In general, Proteobacteria and Cyanobacteria were the most abundant phyla (Figure 1b). Most strikingly, there were clear differences in microbial composition for the communities associated with *Hypotrachyna* and *Cladonia* which had less Cyanobacteria than the communities associated with other lichens. At the class level, Gammaproteobacteria, Alphaproteobacteria, and Nostocophycideae were the most abundant taxa (Figure 1b). Again, there were taxonomic differences at the class level among lichen genera, such as the low abundance observed for Gammaproteobacteria in *Hypotrachyna. Cladonia, Usnea* and *Hypotrachyna* also had very low mean abundance of the Nostocophycideae within their microbiomes, which was more abundant in *Cora*. To obtain an overview of the similarities and differences in taxonomy of the lichen microbiomes sampled, we generated cladograms with the OTU sequences present in all samples from a given lichen genus (Figure 1c). This taxonomy depicts the predominance of taxa from the phylum Proteobacteria in all lichens and the similarity in taxonomic composition of these seven lichen genera at the phylum level, despite differences in the number of identified OTUs, ranging from the lowest number in *Peltigera* to the highest in *Hypotrachyna*.

### Lichens can define microbiomes and share core members

We next analyzed if microbiomes differed based on lichen genus. A hierarchical clustering of lichen sample community composition based on a pairwise Bray-Curtis dissimilarity matrix, indicated that the microbiomes of samples belonging to the same lichen genus were more similar to one another that to those present in different lichen genera (Figure S3). To determine the specific taxa driving these difference we used ALDEx2 (34) software to identify OTUs that were significantly different in abundance among lichen genera. In total we identified 177 OTUs with significant differences as determined by the expected p-value of the Kruskal-Wallace test and the general lineal model-ANOVA (p<0.05). To compare the various lichen microbiomes, we constructed a prevalence matrix based on the presence/absence of OTUs using these 177 differentially abundant taxa. As can be seen in Figure 2a, the microbiomes from samples belonging to the same lichen genus were more similar to one another than to those from other lichens. A Principal Coordinates Analysis (PCoA) also showed that the microbial communities associated with *Cora* lichens, and to a lesser extent with *Cladonia*, clustered close together (Figure 2b), whereas no such clustering was observed when samples were distinguished by growth surface (rock, soil or tree bark) or geographical location (Chingaza vs Nevados Paramo) (Figure S4). A Neighbor-Joining tree constructed with these 177 OTU sequences again revealed a preponderance of taxa from the Proteobacteria, although the relative abundance of taxonomic families within the Proteobacteria varied according to the lichen genus (Figure S5). For example, the Acetobacteraceae family was more abundant in *Cladonia* and *Usnea*, unlike the family Sphingomonadaceae that was more abundant in *Cora* and *Sticta* lichens. Other phyla such as Bacteroidetes, Verrucomicrobia, Cyanobacteria, Acidobacteria and Actinobacteria were also well represented and showed differences in abundance across lichens. Some phyla, like Armatimonadetes, Firmicutes and TM7, were represented by a single OTU. These analyses indicate that the microbial community is predominantly defined by the lichen host rather than by location or growth substrate.

**Figure 2.**
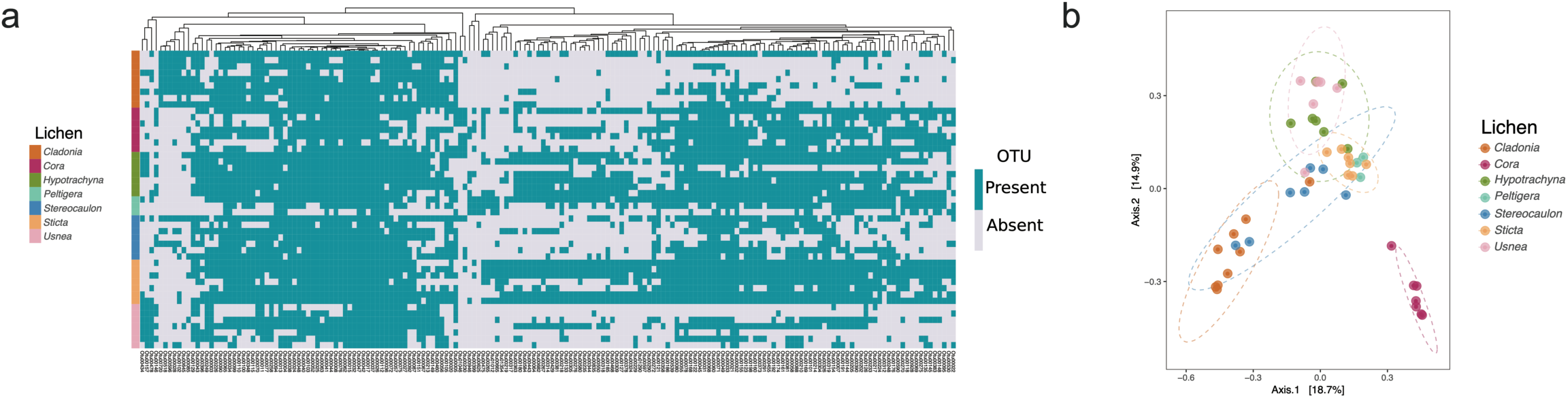
Members of the same lichen genus share similar microbial communities. Data based on the 177 OTUs with significant differences in abundance found by ALDEx2. **a.** Prevalence matrix based on OTU presence/absence. A WPGMA hierarchical clustering method was used to group OTUs on a dendrogram based on a Bray-Curtis dissimilarity matrix. **b.** Principal Coordinates Analysis (PCoA) based on the Bray-Curtis index shows microbiomes of lichens *Cora* and *Cladonia* differentiated from other lichens.

To detect if lichens harbored a core microbiome, shared OTUs were identified by registering both the presence (prevalence) and the total counts (abundance) for each of the originally identified 20,174 OTUs in all samples. An OTU was considered to be part of the core microbiome if it was present in at least 90% of samples (≥ 90%) (Figure 3a). OTUs with a prevalence < 25% were cataloged as *peripheral*, taxa that might be an extension of the environment or substrate on which the lichen grows. OTUs with a prevalence between ≥ 25 and < 90% represent *pan* taxa that might be occasionally present in lichens but are not widely distributed across samples and lichen genera. Sixteen OTUs were shared among all lichens sampled (Figure 3a). Their abundances ranged from 2,777 to 29,245 counts per OTU. Core OTUs corresponded to Proteobacteria (eleven OTUs), Acidobacteria (four OTUs), and Cyanobacteria (one OTU). The eleven Proteobacteria OTUs belonged to three orders, Rhizobiales, Rhodospirillales and Sphingomonadales, while the Acidobacteria OTUs corresponded only to the order Acidobacteriales. The Cyanobacteria OTU remained unclassified according to the taxonomic assignment with the Greengenes database. While these sixteen core taxa represented a minor part of the total number of OTUs (Figure 3a) they were among the most abundant in the data set. However, their abundance was variable among the different lichen genera, as can be seen for the Cyanobacteria OTU (Figure S6). Thus, these diverse lichens appear to harbor a small core microbiome.

**Figure 3.**
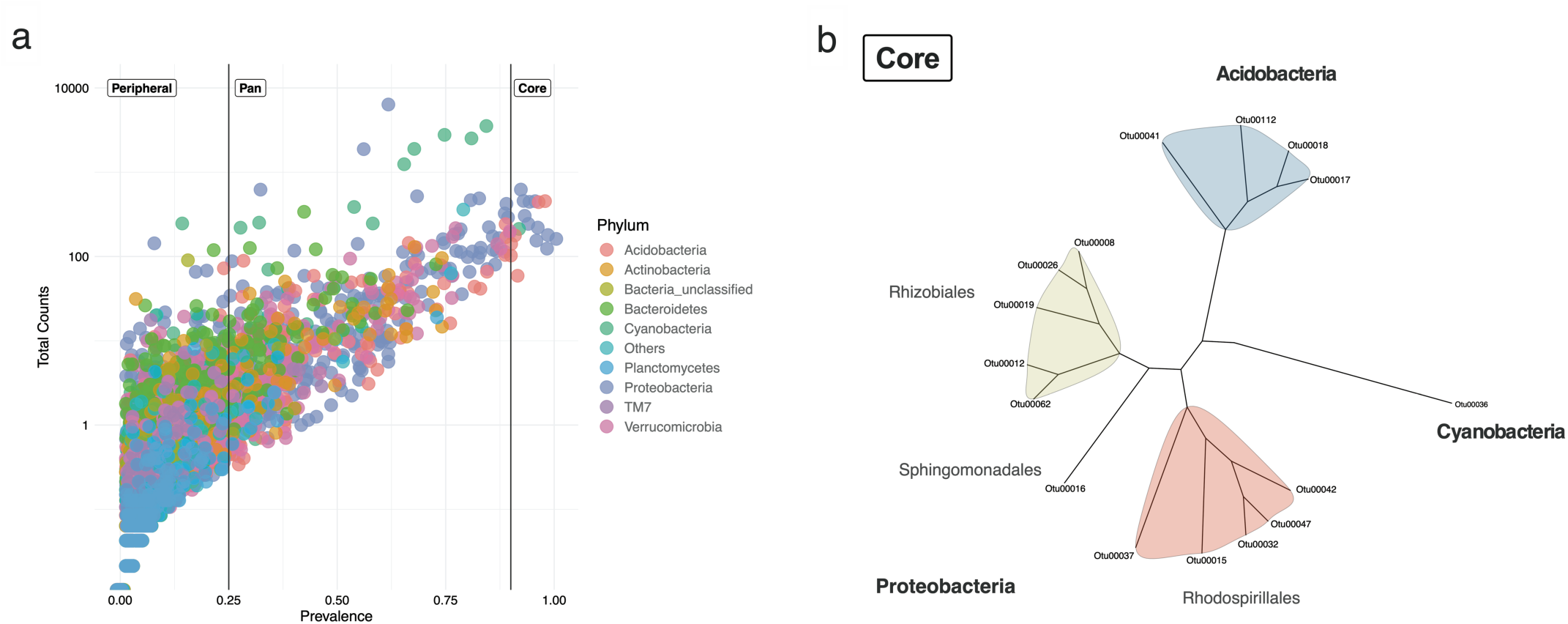
Lichen core microbiome of samples from two Paramos. **a.** Prevalence of the total 20,174 OTUs across the 47 lichen samples and their total number of observations (counts). Taxa were defined as Core (prevalence ≥ 0.9), Pan (prevalence ≥ 0.25 and < 0.9) and Peripheral (prevalence < 0.25). OTUs are colored by phylum. **b.** Neighbor-Joining tree of the lichen core microbiome.

To see if these sixteen core taxa were also present in lichens from different geographical sites, we aligned our sequences against 16S rRNA gene sequences available from the NCBI database (Identity values >98%). Of 34 sequences identified, eleven corresponded to uncultured bacteria from lichens (Figure S7). However only in one case the lichen species was identified (*Ramalina pollinaria,* GenBank-ID MG996731.1) (Table S3). Most of the 34 sequences retrieved from NCBI corresponded to samples found in cold environments such as Glaciers and Tundra, which have environmental conditions comparable to those of the Paramo. These results suggest that the OTUs of our core microbiome are similar to bacteria found in lichens described in other studies.

### Paramo lichens are a rich source of antimicrobials

In order to investigate if the studied lichens harbored bacteria that produce antimicrobials, the original 57 lichens collected in both parks were processed by plating on multiple media. 122 isolates were obtained from 37 samples belonging to eleven lichen genera (Table S4). Of these isolated strains, 112 were Bacteria and 10 were Fungi, based on PCR amplification of the 16S rRNA gene and ITS region, respectively. Approximately 62% of these isolates (n=76) were obtained from lichens collected in Chingaza, while 38% (n=46) were from Nevados. All isolates were tested for antimicrobial activity using a double agar layer assay against seven pathogens (*Staphylococcus aureus, Acinetobacter baumannii, Pseudomonas aeruginosa, Candida albicans, Salmonella enterica, Escherichia coli* and *Klebsiella pneumoniae*). 28% of the bacterial strains (n=32) and 30% of fungi (n=3) displayed antimicrobial activity against at least one of the pathogens tested, the majority of which (n=26) were recovered from Chingaza lichens. We found antimicrobial activity against all seven pathogens, but the most detected activities were against *A. baumannii, S. aureus*, and *C. albicans*. Of the 35 isolates showing antimicrobial activity, 21 strains were active against multiple pathogens. Additionally, two bacteria isolated from lichens *Psoroma* and *Yoshimuriella* exhibited activity against a multi-drug-resistant *Klebsiella pneumoniae* strain cataloged in our laboratory as resistant to β-lactams (Cephalosporins, β-lactamase Inhibitors, Carbapenems, Monobactams), Amikacin, Ciprofloxacin and Meropenem based on Antimicrobial Susceptibility Standards values (35).

## Discussion

Despite the importance of lichens for ecosystems, there is still limited understanding of their biology and, especially, of the assembly and function of the associated microbial communities. The pervasiveness of microbial communities associated with lichens suggest that at least some of these microbes may be more than transient associates in these symbiotic structures. Here we describe the microbiomes of lichens from the Andean Paramo ecosystems, high mountain habitats that harbor endemic species and are an important reservoir of lichen diversity. Colombia is known to harbor at least 10% of the described lichen species described in the world (36, 37), likely a low-end estimate (31). This study extends previous observations by characterizing and comparing microbiomes present in seven lichen genera, *Cora, Hypotrachyna, Cladonia, Usnea, Sticta, Stereocaulon* and *Peltigera* and, to our knowledge the first assessment of the microbiome composition of *Cora* and *Hypotrachyna* lichens. Here we identified both transient members, expected due to location and exposure to variable environmental conditions, and more permanently associated taxa that indicate a tight relationship and can orient studies aimed at understanding how these microbes contribute to lichen function and ecology.

The 16S rRNA gene analyses of these Paramo lichen microbiomes showed diverse and complex microbial taxonomic profiles that varied among samples, as has been observed in other studies (38). The seven lichen genera sampled harbored microbiomes composed mainly of the phyla Proteobacteria, Acidobacteria, Verrucomicrobia, Bacteroidetes and Actinobacteria, consistent with previous reports (39, 40). There was, however, variation in the relative abundance of these groups across the various lichens and among individual samples of the same lichen genus, an indication of the heterogeneity that can be expected in environmental microbiomes. The observed variation in community structures, particularly within a given genus, could be due to the fact that not all lichens were classified to the species level and therefore the samples from a given genus could include multiple species. In fact, some of the genera sampled here are considered to be among the largest in terms of the number of lichenized species, with *Cladonia* containing the most known species among the lichens studied (41).

Despite the observed variability, lichen microbiomes showed a high abundance of Proteobacteria, as has been documented in other studies (30, 42). Members of this phylum are thought to play important roles in lichen symbioses by providing nutrients, mobilizing iron and phosphate, and fixing nitrogen (24, 43). Among the Proteobacteria, Gammaproteobacteria and Alphaproteobacteria were the most abundant groups. Unlike other studies of lichen microbiomes, the Gammaproteobacteria were more abundant than the Alphaproteobacteria in many of our samples. Alphaproteobacteria has been consistently reported as an abundant member of the bacterial microbiome in lichens (39, 40, 42, 44). The detection of Gammaproteobacteria and their predominance over Alphaproteobacteria in our dataset, differs from previous studies and could be due to the fact that here we sampled different lichen genera from a novel geographic region (44, 45). It is also possible that this discrepancy is due to methodological differences such as sample collection, processing, and sequencing and analysis platforms, all of which can have an effect on the resulting community profiles and subsequent inferences.

Taxonomic profiling revealed Cyanobacteria as abundant members of all microbiomes. In fact, Cyanobacteria were present in *Cladonia, Hypotrachyna* and *Usnea*, which are considered to be chlorolichens (46-49), that is to say, lichens with a green alga as its major or unique photobiont (50). Tripartite lichens, which have both an algal and a cyanobacterial photobiont are known to make up a small number of lichens (about 3-4%) and introduce greater complexity to these structures since both photobionts can contribute to photosynthesis (51, 52). While the presence of abundant Cyanobacteria could suggest an important role in these chlorolichens, such as nitrogen fixation and/or photosynthesis, further analyses would have to be done to confirm their precise functions. Given the limitation of Illumina sequencing, which provides information for only ∼300bp of the V3-V4 portion of the 16S rRNA gene, these Cyanobacterial taxa could not be classified at higher phylogenetic levels with the available databases (Figure S8). Additional metagenomic sequencing or the isolation of these microbial members would be needed to further determine if these are novel cyanobacterial species and to assess their possible roles within lichens.

Several studies have identified patterns in the structure of lichen microbial communities (30, 39, 45, 53). Specific bacterial taxa have been associated with some lichens (54), as well as predominance of particular groups, such as the Alphaproteobacteria in *Cladonia arbuscula* (42). These differences are thought to be driven by biotic and abiotic factors, of which the photobiont and large-scale geographical distance apparently determine the composition in different lichen types (29). It has also been suggested that a lichen’s secondary metabolite production could drive microbiome structure (29, 55, 56). In this work, we used a differential abundance analysis approach (ALDEx2) to identify OTUs that varied in abundance among samples. ALDEx2 takes into account the compositional nature of microbiome data (57), which means that the limited number of sequences obtained in any sequencing platform do not necessarily represent the number of sequences present in a given sample. This pipeline considers sample variation, which can be due to technical variations such as library preparation and sequencing output, to identify taxa that are significantly different between groups, reducing the false discovery rate frequently associated with other standard approaches for high-throughput sequencing data (34, 57, 58). This strategy also removes biases associated with standard data analysis that frequently defines bacterial community patterns based mostly on abundant taxa (11), and can overlook rare OTUs or low abundant taxa that may be important for host function (59). This bias is evident in a variety of ecosystems where rare taxa have been seen to be essential for the dynamics of microbial communities (60, 61).

By using this differential abundance analysis, we identified a set of 177 OTUs with significant differences, from a total of 20,174 OTUs, that indicated that these microbiomes were not defined by geographical location or growth substrate (foliose, fruticose, or crustose), as reported in some cases (30). In our work, lichen microbiomes appeared to be driven by the lichen genus, for the case of *Cora* and to a lesser extent for *Cladonia* samples. Interestingly, *Cora* is the only lichen sampled here that is known to have a Basidiomycete mycobiont (41), suggesting that the fungal host could be important in shaping this microbiome. For *Cladonia* lichens, the community clustering was not as evident but might be further examined by identifying if lichen species play a role in defining community structure (29), something that could not be done given that we did not classify the lichens to the species level (45). The 177 significantly different OTUs mainly belonged to the class Alphaproteobacteria (Figure S5). However, the relative abundance of Alphaproteobacteria varied across the seven lichen genera. Interestingly, some taxa from the family Acetobacteraceae were more abundant in lichens *Cladonia* and *Usnea*, while the Sphingomonadaceae taxa were more prominent in lichens *Cora* and *Sticta*. Some authors have hypothesized that this variation of Alphaproteobacteria abundance might depend on the type of lichen photobiont (29), with the order Rhodospirillales dominating in chlorolichens and Sphingomonadales in cyanolichens. However further analyses to determine the type of photobiont in our samples are needed in order to explain the variation of Alphaproteobacteria taxa.

A core microbiome of 16 OTUs was identified across the 47 lichens sampled from two distant Paramos. The limited number of shared OTUs reflects the complexity and diversity of these lichen microbiomes and the fact that seven different genera, and possibly many uncharacterized species, were analyzed. Some of these core taxa were found to be similar to 16S rRNA gene sequences found in other lichens from distant sites including extreme environments such as the Arctic. Further sampling and deeper sequencing efforts might help to determine if this core is in fact conserved in other lichens. In contrast, the same analysis within a more tightly defined phylogenetic group, such as a single species, could identify a more robust core community. The lichen core microbiome included representatives of three phyla, Proteobacteria, Acidobacteria and Cyanobacteria. The Proteobacteria core members belonged to the Alphaproteobacteria class, which functional omics studies have highlighted as essential for nutrient supply and lichen growth (39, 42, 54, 62). These Alphaproteobacteria were represented by the orders Rhizobiales, Rhodospirillales and Sphingomonadales that have been previously reported as crucial for the maintenance of lichens (54, 62). Finally, our core microbiome indicated the presence of a single cyanobacterial OTU, even though there were other highly abundant Cyanobacteria in >60% of our lichen samples (*Pan* microbiome). Previous reports have shown that cyanobacterial symbionts can be shared among different lichen types (63), while in other cases the lichen mycobiont might be strongly selective in the choice of cyanobiont (64, 65). Future studies focused on cyanobacterial specificity within lichen microbiomes could disentangle the roles that these taxa are playing within lichen thalli.

Soil bacteria have traditionally been the major source of antimicrobials (66, 67), but most of these compounds are derived from relatively few culturable microbial taxa (68). With the rapid and widespread increase of multi-drug-resistant bacteria, there is a pressing need for new antimicrobials (69) that has prompted exploration of different ecosystems. Bacteria producing bioactive compounds have been isolated from some lichens such as *Lobaria* (28), and *Cladonia* (70) and even from marine lichens (71). These bacteria belong mainly to the phylum Actinobacteria, a group well-known for its biosynthetic capacity and antimicrobial production (72, 73). Members of this phylum have been consistently reported as members of lichen microbiomes (39, 53, 74), and in situ analyses of *C. arbuscula* have shown that these bacteria are located within the thallus structure (42). Our lichen microbiomes had a high abundance of actinobacterial taxa, and antimicrobial screening showed that lichen bacterial isolates produced molecules active against diverse microorganisms, including the multi-drug-resistant pathogen *K. pneumoniae*. Taken together, these results suggest that lichen microbiomes from underexplored ecosystems such as the Paramo, could be an important source of novel bacteria and antimicrobials. These antimicrobial-producing bacteria could be crucial for the defense of lichen thalli against pathogens or for the maintenance of microbial community balance within the symbiosis. In addition to further analyses of potential bioactive compounds, metagenomic studies of our lichen isolates should help to identify bacterial species, biosynthetic gene clusters and their metabolic potential.

## Conclusions

Here we described the microbiomes of seven lichen genera (*Usnea, Cladonia, Peltigera, Stereocaulon, Sticta, Cora* and *Hypotrachyna*), including the first description of the bacterial communities from *Cora* and *Hypotrachyna* lichens, and the presence of a core lichen microbiome. These Paramo lichen microbiomes were dominated by the phyla Proteobacteria, Cyanobacteria, Acidobacteria, Verrucomicrobia, Bacteroidetes and Actinobacteria. These microbiomes varied among lichens and were distinguished based on host identity rather than location or growth substrate. Importantly, we found a core community of sixteen OTUs present in all samples. The core community was composed of members from only three phyla, Proteobacteria, Acidobacteria and Cyanobacteria, suggesting that there is high selectivity regarding which bacteria can establish close associations across all lichens. Microbes isolated from these lichens produced antifungal and antibacterial compounds which suggests that these ecosystems could be further probed as a source of natural products.

## Methods

### Sampling and sample processing

Samples were taken at Los Nevados and Chingaza National Natural Parks in Colombia at altitudes ranging from 3,600 to 4,160 m.a.s.l using sterile forceps and immediately placed in sterile plastic bags at environmental temperature. Several individual thalli were taken in order to have representative samples of different lichen genera in both localities. Samples were processed within 48 hours of collection. Metadata such as GPS location and types of substrate where the lichen was collected (Corticolous: Wood, Terricolous: Soil and Saxicolous: Rock) were taken (Table S1). Lichen morphological identification at genus level was carried out through herbarium specimen comparison.

Genomic DNA of lichens was extracted using the PowerSoil DNA Isolation Kit (Qiagen) with some modifications: 20mg of lichen were homogenized in a FastPrep-24 (MP Biomedicals) for two 20 second cycles at 4 m/s, and then processed according to the manufacturer instructions. The 16S rRNA gene V3-V4 region was amplified with primers V3F (5’-CCTACGGGAGGCAGCAG-3’) and V4R (5’-GGACTACHVGGGTWTCTAAT-3’) with barcoded Illumina adapters as describe in the standard procedures of the Earth Microbiome Project (http://www.earthmicrobiome.org/protocols-and-standards/16s/). Blank controls were also included in amplification for quality assurance. Each 20μL PCR reaction was prepared with 4μL 5x HOT FIREPol master mix (Solis BioDyne), 2μL of each primer (10μM), 2μL of sample DNA and 12μL PCR-grade water. The amplicons were pooled in equimolar concentrations using SequalPrep plate normalization kit (Invitrogen) and then purified with AMPure XP beads (Beckman Coulter). Amplicons were sequenced on the Illumina MiSeq platform at the DNA Services Facility at the Microbiology Department, Harvard Medical School in Boston, USA.

### Analyses of sequence data

Illumina reads were quality checked with FastQC and edited with Trimmomatic (75) to remove adapter and low-quality sequences that included reads with ambiguous nucleotides (Q value<25) and short reads (<200bp). Edited reads were processed in Mothur (v1.40) (76), by first removing sequences longer than 430pb (screen.seqs: maxambig=0, maxlength=430). Files were reduced to non-identical sequences (unique.seqs and count.seqs) to minimize computational effort. Non-redundant sequences were aligned (align.seqs) to a trimmed SILVA (v132) bacteria database (pcr.seqs: start=7697, end=23444, keepdots=F) provided by Mothur (77). Only sequences that were aligned to the expected position were kept (screen.seqs start=2, end=15747, maxhomop=8; filter.seqs: vertical=T, trump=.). Aligned sequences were again reduced to non-redundant sequences and de-noised (unique.seq; pre.cluster), checked for chimeras using the VSEARCH algorithm (chimera.vsearch: dereplicate=t), which were then filtered out (remove.seqs). Sequences were classified (classify.seqs) based on the Greengenes database provided by Mothur (78). Possible undesirable misclassified lineages were removed (remove.lineage taxon=Chloroplast-Mitochondria-unknown-Archaea-Eukarya). Sequences were then clustered (cluster.split: splitmethod=classify, taxlevel=4, cutoff=0.03) and converted to shared file format (make.shared: label=0.03) assigning taxonomy to each OTU (classify.otu: label=0.03, relabund=t). For alpha-diversity analysis reads were normalized to 20,623. Representative sequences of OTUs were retrieved based on the distance among the clustered sequences (get.oturep). The non-normalized shared file with OTU counts was used for differential abundance analysis in beta-diversity with ALDEx2 (79).

### Diversity comparisons and statistical analyses

Diversity within samples (alpha-diversity) was analyzed with the Shannon-Weaver (80) and Simpson Index (81). Richness of microbial communities was assessed based on the observed number of OTUs and the rarefaction curves using the R package Phyloseq (82). Multiple comparisons of richness and diversity measures were performed by one-way ANOVA, including Tukey’s (equal SD) or Tamhane T2 (non-equal SD) corrections. P values of <0.05 were considered to be statistically significant. Microbial community comparisons (beta-diversity) were first assessed with a similarity tree of samples based on the Bray-Curtis distance similarity matrix and the WPGMA hierarchical clustering method. We used ALDEx2 analysis (ANOVA-Like Differential Expression tool for compositional data) (83) to find OTUs that define the differences between lichen microbiomes. ALDEx2 R package decomposes sample-to-sample variation into four parts (within-condition variation, between-condition variation, sampling variation, and general unexplained error) using Monte-Carlo sampling from a Dirichlet distribution (aldex.clr: denom=“all”) (84, 85). The statistical significance of each OTUs was determined by the general lineal model and Kruskal-Wallis Test (aldex.kw) for one-way ANOVA to determine OTUs significantly different for the seven lichen genera under study. The significantly differentially abundant OTUs were used to generate a Principal Coordinate Analysis (PCoA) based on the Bray-Curtis index and a prevalence matrix based on presence/absence. A Neighbor-Joining tree with differentially abundant OTUs and their abundances was built with OTU sequences aligned by an iterative refinement method (FFT-NS-i) (86, 87).

To display the taxonomy of OTUs present in each lichen microbiome, sequences were aligned in MAFFT v.7 with default settings (87), and the cladogram for each microbiome was constructed using the average linkage method (UPGMA) (88).

### Core microbiome

OTU prevalence (20,174 OTUs) was calculated based on the count mean of each OTU in every sample and cataloged as core (prevalence ≥ 0.9), *pan* (prevalence ≥ 0.25 and <0.9) or *peripheral* (<0.25). Core OTU sequences were aligned by an iterative refinement method (FFT-NS-i) and clustered by Neighbor-Joining (Jukes-Cantor Model) on MAFFT v.7 (87). Core OTU relative abundances (CLR-transformed) in each lichen genus were displayed on a violin plot from Prism8. Core OTUs sequences were aligned to sequences in NCBI using Blastn optimized for highly similar sequences. Reference sequences were chosen based on >98% identity value. Both reference and core sequences were aligned and clustered with the same parameters mentioned above.

### Bacterial Isolation and Antimicrobial screen

Lichens were briefly washed with sterile water to remove sediment and loosely attached microorganisms (71, 89). Samples were aseptically divided into small pieces (∼0.5 cm) using sterile scalpels. The pieces were homogenized in phosphate saline solution with glass beads (4mm-Marienfield) using a vortex. 100μL of 10-fold serial dilutions in phosphate saline solution were plated on four culture media: a) Actinomycete Isolation Agar (AIA, Difco, BD), b) International Streptomyces Project medium-2 (ISP2), both supplemented with nalidixic acid (150mg/L) and nystatin (50mg/L) (67, 89), c) Gause Synthetic medium, and d) Gause Oligotrophic supplemented with potassium dichromate (80mg/L) (90).

Plates were incubated at room temperature (∼25 °C) until no more new colonies appeared (up to 20 days). Colonies were isolated and purified based on morphological characteristics [color, surface (smooth or rough), shape (circular, filamentous, irregular or punctiform) and edge format (regular or irregular)]. DNA was extracted from purified colonies using the phenol chloroform extraction method with the following modifications: glass beads (0.4mm) were used to lyse cells in a FastPrep-24 homogenizer (MP Biomedicals) with two cycles of 20 seconds at 4 m/s, adding 500μL Tris-HCl buffer p.H 8.0, 200μL NaCl 2.8M and 34μL SDS 0.8% in 2mL tubes. Bacterial strains were stored in the same medium in which they were isolated with 20% glycerol at −80°C.

Bacterial DNA was used to amplify the 16S rRNA using 27F and 1492R universal primers (91): 27F 5’-AGAGTTTGATCCTGGCTCAG-3’ and 1492R: 5’-ACGGTTACCTTGTTACGACTT-3’. Each 25μL PCR reaction contained 12.5μL CorpoGen PCR Master mix, 0.5μL of each primer (25μM), 9.5μL PCR-grade water. PCR amplification was done by 3 min denaturation at 94°C; 35 cycles of 30s at 94°C, 45s at 55°C and 60s at 72°C; and 6 min elongation at 72°C. Isolates negative for 16S rRNA gene were corroborated as fungi by amplifying the ITS region with primers ITS5 and ITS4 (92).

Antimicrobial screening was performed using the double agar layer assay (93), against seven pathogens of medical importance: *S. enterica, E. coli, K. pneumoniae, P. aeruginosa, A. baumannii, S. aureus* and *C. albicans*. Isolated strains were grown on solid medium for 10 days, covered with Mueller-Hinton agar containing 100 μL of an overnight culture of each pathogen (94), and incubated at 37 °C for 24 hours. Strains that displayed a growth inhibition halo of the tester pathogens were considered as antimicrobial producers.

## Data availability

Sequence data of lichen microbiomes are available in NCBI under accession number PRJNA558995. OTUs and taxonomy tables together with the figure scripts are available on GitHub: https://github.com/mariaasierra/Lichen_Microbiome

## Author contributions

MAS carried out sampling, laboratory work, data analysis and manuscript writing. TS, DD and CM helped with data and statistical analyses and manuscript editing. GP helped with sequencing. RK helped with sampling and analysis of data. BM carry out identification of lichens. MMZ conceived the study, supervised work, helped with sampling and with writing of the manuscript. All authors provided input on the manuscript, read and agreed to the contents of the final version.

## Acknowledgements

The authors wish to thank Adan Ramirez-Rojas and Angela Cantillo for their help in sampling and processing DNA. This work was financed by Colciencias (project No. 639676359556) and was carried out under permits for sample collection in Colombia (ANLA permit 1484 and National Parks permit No. 13-2017).

## Competing interests

The authors declare that they have no competing interests

## Notes

https://github.com/mariaasierra/Lichen_Microbiome

## Bibliography

1. Gilbert SF, Sapp J, Tauber AI. A symbiotic view of life: we have never been individuals. Q Rev Biol. 2012;87(4):325–41.

2. de Zelicourt A, Al-Yousif M, Hirt H. Rhizosphere microbes as essential partners for plant stress tolerance. Molecular plant. 2013;6(2):242–5.

3. Douglas AE. The microbial dimension in insect nutritional ecology. Functional Ecology. 2009;23(1):38–47.

4. Ceja-Navarro JA, Karaoz U, Bill M, Hao Z, White RA, Arellano A, et al. Gut anatomical properties and microbial functional assembly promote lignocellulose deconstruction and colony subsistence of a wood-feeding beetle. Nature microbiology. 2019:1.

5. Berg M, Koskella B. Nutrient-and dose-dependent microbiome-mediated protection against a plant pathogen. Current Biology. 2018;28(15):2487-92. e3.

6. Berendsen RL, Pieterse CM, Bakker PA. The rhizosphere microbiome and plant health. Trends in plant science. 2012;17(8):478–86.

7. Thomas T, Moitinho-Silva L, Lurgi M, Björk JR, Easson C, Astudillo-García C, et al. Diversity, structure and convergent evolution of the global sponge microbiome. Nature communications. 2016;7:11870.

8. Shapira M. Gut microbiotas and host evolution: scaling up symbiosis. Trends in ecology & evolution. 2016;31(7):539–49.

9. Hacquard S, Garrido-Oter R, González A, Spaepen S, Ackermann G, Lebeis S, et al. Microbiota and host nutrition across plant and animal kingdoms. Cell host & microbe. 2015;17(5):603–16.

10. Ochman H, Worobey M, Kuo C-H, Ndjango J-BN, Peeters M, Hahn BH, et al. Evolutionary relationships of wild hominids recapitulated by gut microbial communities. PLoS biology. 2010;8(11):e1000546.

11. Shade A, Handelsman J. Beyond the Venn diagram: the hunt for a core microbiome. Environmental microbiology. 2012;14(1):4–12.

12. T DA, Krause L, Bridge T, Torda G, Raina JB, Zakrzewski M, et al. The coral core microbiome identifies rare bacterial taxa as ubiquitous endosymbionts. ISME J. 2015;9(10):2261–74.

13. Sanders JG, Powell S, Kronauer DJ, Vasconcelos HL, Frederickson ME, Pierce NE. Stability and phylogenetic correlation in gut microbiota: lessons from ants and apes. Molecular Ecology. 2014;23(6):1268–83.

14. Yeoh YK, Dennis PG, Paungfoo-Lonhienne C, Weber L, Brackin R, Ragan MA, et al. Evolutionary conservation of a core root microbiome across plant phyla along a tropical soil chronosequence. Nature communications. 2017;8(1):215.

15. Ley RE, Hamady M, Lozupone C, Turnbaugh PJ, Ramey RR, Bircher JS, et al. Evolution of mammals and their gut microbes. Science. 2008;320(5883):1647–51.

16. Yuan X, Xiao S, Taylor TN. Lichen-like symbiosis 600 million years ago. Science. 2005;308(5724):1017–20.

17. Nash TH. Lichen Biology: Cambridge: Cambridge University Press.; 2008.

18. Seneviratne G, Indrasena I. Nitrogen fixation in lichens is important for improved rock weathering. Journal of biosciences. 2006;31(5):639–43.

19. Nieboer E, Richardson D, Tomassini F. Mineral uptake and release by lichens: an overview. Bryologist. 1978:226–46.

20. Thomas H. Nash I. Lichen Biology. 2 ed. Cambridge: Cambridge University Press; 2008.

21. Shimizu A. Community structure of lichens in the volcanic highlands of Mt. Tokachi, Hokkaido, Japan. Bryologist. 2004:141–51.

22. Garvie LA, Knauth LP, Bungartz F, Klonowski S, Nash TH. Life in extreme environments: survival strategy of the endolithic desert lichen Verrucaria rubrocincta. Naturwissenschaften. 2008;95(8):705–12.

23. Cardinale M, Puglia AM, Grube M. Molecular analysis of lichen-associated bacterial communities. FEMS Microbiology Ecology. 2006;57(3):484–95.

24. Grube M, Cernava T, Soh J, Fuchs S, Aschenbrenner I, Lassek C, et al. Exploring functional contexts of symbiotic sustain within lichen-associated bacteria by comparative omics. The ISME Journal. 2015;9(2):412–24.

25. Cernava T, Erlacher A, Aschenbrenner IA, Krug L, Lassek C, Riedel K, et al. Deciphering functional diversification within the lichen microbiota by meta-omics. Microbiome. 2017;5(1):82.

26. Spribille T, Tuovinen V, Resl P, Vanderpool D, Wolinski H, Aime MC, et al. Basidiomycete yeasts in the cortex of ascomycete macrolichens. Science. 2016;353(6298):488–92.

27. Davies J, Davies D. Origins and evolution of antibiotic resistance. Microbiology and molecular biology molecular. 2010;74(3):417–33.

28. Cernava T, Aschenbrenner IA, Grube M, Liebminger S, Berg G. A novel assay for the detection of bioactive volatiles evaluated by screening of lichen-associated bacteria. Frontiers in microbiology. 2015;6:398.

29. Hodkinson BP, Gottel NR, Schadt CW, Lutzoni F. Photoautotrophic symbiont and geography are major factors affecting highly structured and diverse bacterial communities in the lichen microbiome. Environmental microbiology. 2012;14(1):147–61.

30. Aschenbrenner IA, Cardinale M, Berg G, Grube M. Microbial cargo: do bacteria on symbiotic propagules reinforce the microbiome of lichens? Environmental microbiology. 2014;16(12):3743–52.

31. Lücking R, Dal-Forno M, Sikaroodi M, Gillevet PM, Bungartz F, Moncada B, et al. A single macrolichen constitutes hundreds of unrecognized species. Proceedings of the National Academy of Sciences. 2014;111(30):11091–6.

32. Myers N, Mittermeier RA, Mittermeier CG, da Fonseca GAB, Kent J. Biodiversity hotspots for conservation priorities. Nature. 2000;403(6772):853–8.

33. Sabater S, González-Trujillo JD, Elosegi A, Rondón JCD. Colombian ecosystems at the crossroad after the new peace deal. Biodiversity and Conservation. 2017;26(14):3505–7.

34. Fernandes AD, Macklaim JM, Linn TG, Reid G, Gloor GB. ANOVA-like differential expression (ALDEx) analysis for mixed population RNA-Seq. PLoS One. 2013;8(7):e67019.

35. Clinical and Laboratory Standards Institute (CLSI). Performance Standards for Antimicrobial Susceptibility Testing. 29th ed. CLSI supplement M100 (ISBN 978-1-68440-032-4 [Print]; ISBN 978-1-68440-033-1 [Electronic]). Clinical and Laboratory Standards Institute, 950 West Valley Road, Suite 2500, Wayne, Pennsylvania 19087 USA, 2019.

36. Bernal R, S.R. Gradstein & M. Celis (eds.). Catálogo de plantas y líquenes de Colombia. Instituto de Ciencias Naturales, Universidad Nacional de Colombia, Bogotá.2015 [

37. Gonzalez-Salazar MA, Venturini M, Poganietz W-R, Finkenrath M, L.V. Leal MR. Combining an accelerated deployment of bioenergy and land use strategies: Review and insights for a post-conflict scenario in Colombia. Renewable and Sustainable Energy Reviews. 2017;73:159–77.

38. Aschenbrenner IA, Cernava T, Berg G, Grube M. Understanding microbial multi-species symbioses. Frontiers in Microbiology. 2016;7(FEB):1–9.

39. Bates ST, Cropsey GWG, Caporaso JG, Knight R, Fierer N. Bacterial communities associated with the lichen symbiosis. Applied and Environmental Microbiology. 2011;77(4):1309–14.

40. Grube M, Berg G. Microbial consortia of bacteria and fungi with focus on the lichen symbiosis. Fungal biology reviews. 2009;23(3):72–85.

41. Lücking R, Dal Forno M, Moncada B, Coca LF, Vargas-Mendoza LY, Aptroot A, et al. Turbo-taxonomy to assemble a megadiverse lichen genus: seventy new species of Cora (Basidiomycota: Agaricales: Hygrophoraceae), honouring David Leslie Hawksworth’s seventieth birthday. Fungal Diversity. 2017;84(1):139–207.

42. Cardinale M, Vieira de Castro J, Jr., Muller H, Berg G, Grube M. In situ analysis of the bacterial community associated with the reindeer lichen Cladonia arbuscula reveals predominance of Alphaproteobacteria. FEMS Microbiol Ecol. 2008;66(1):63–71.

43. Sigurbjörnsdóttir MA, Vilhelmsson O. Selective isolation of potentially phosphate-mobilizing, biosurfactant-producing and biodegradative bacteria associated with a sub-Arctic, terricolous lichen, Peltigera membranacea. FEMS microbiology ecology. 2016;92(6).

44. Mushegian AA, Peterson CN, Baker CC, Pringle A. Bacterial diversity across individual lichens. Applied and environmental microbiology. 2011;77(12):4249–52.

45. Grube M, Cardinale M, de Castro JV, Müller H, Berg G. Species-specific structural and functional diversity of bacterial communities in lichen symbioses. The ISME Journal. 2009;3(9):1105–15.

46. Anna V, Dymytrova L, Rai H, Upreti DK. Photobiont diversity of soil crust lichens along substrate ecology and altitudinal gradients in Himalayas: a case study from Garhwal Himalaya. Terricolous lichens in India: Springer; 2014. p. 73–87.

47. Ahmadjian V. The lichen symbiosis: John Wiley & Sons; 1993.

48. Skaloud P, Peksa O. Evolutionary inferences based on ITS rDNA and actin sequences reveal extensive diversity of the common lichen alga Asterochloris (Trebouxiophyceae, Chlorophyta). Molecular Phylogenetics and Evolution. 2010;54(1):36–46.

49. Rafat A, Ridgway HJ, Cruickshank RH, Buckley HL. Isolation and co-culturing of symbionts in the genus Usnea. Symbiosis. 2015;66(3):123–32.

50. Lange OL, Wagenitz G. Vernon Ahmadjian introduced the term ‘chlorolichen’. The Lichenologist. 2004;36(2):171-.

51. Henskens FL, Green TA, Wilkins A. Cyanolichens can have both cyanobacteria and green algae in a common layer as major contributors to photosynthesis. Annals of botany. 2012;110(3):555–63.

52. Rikkinen J. Cyanobacteria in terrestrial symbiotic systems. Modern Topics in the Phototrophic Prokaryotes: Springer; 2017. p. 243–94.

53. Bjelland T, Grube M, Hoem S, Jorgensen SL, Daae FL, Thorseth IH, et al. Microbial metacommunities in the lichen – rock habitat. 2011;3:434–42.

54. Hodkinson BP, Lutzoni F. A microbiotic survey of lichen-associated bacteria reveals a new lineage from the Rhizobiales. Symbiosis. 2009;49(2):163–80.

55. Leiva D, Clavero-León C, Carú M, Orlando J. Intrinsic factors of Peltigera lichens influence the structure of the associated soil bacterial microbiota. FEMS microbiology ecology. 2016;92(11):fiw178.

56. Cernava T, Müller H, Aschenbrenner IA, Grube M, Berg G. Analyzing the antagonistic potential of the lichen microbiome against pathogens by bridging metagenomic with culture studies. Frontiers in microbiology. 2015;6:620-.

57. Gloor GB, Macklaim JM, Pawlowsky-Glahn V, Egozcue JJ. Microbiome datasets are compositional: and this is not optional. Frontiers in microbiology. 2017;8:2224.

58. Gloor GB, Reid G. Compositional analysis: a valid approach to analyze microbiome high-throughput sequencing data. Canadian journal of microbiology. 2016;62(8):692–703.

59. Reveillaud J, Maignien L, Eren AM, Huber JA, Apprill A, Sogin ML, et al. Host-specificity among abundant and rare taxa in the sponge microbiome. The ISME journal. 2014;8(6):1198.

60. Shade A, Jones SE, Caporaso JG, Handelsman J, Knight R, Fierer N, et al. Conditionally rare taxa disproportionately contribute to temporal changes in microbial diversity. MBio. 2014;5(4):e01371–14.

61. Dohrmann AB, Küting M, Jünemann S, Jaenicke S, Schlüter A, Tebbe CC. Importance of rare taxa for bacterial diversity in the rhizosphere of Bt-and conventional maize varieties. The ISME journal. 2013;7(1):37.

62. Erlacher A, Cernava T, Cardinale M, Soh J, Sensen CW, Grube M, et al. Rhizobiales as functional and endosymbiontic members in the lichen symbiosis of Lobaria pulmonaria L. Front Microbiol. 2015;6:53.

63. Rikkinen J, Oksanen I, Lohtander K. Lichen guilds share related cyanobacterial symbionts. Science. 2002;297(5580):357-.

64. O’Brien HE, Miadlikowska J, Lutzoni F. Assessing host specialization in symbiotic cyanobacteria associated with four closely related species of the lichen fungus Peltigera. European journal of phycology. 2005;40(4):363–78.

65. Rikkinen J. Molecular studies on cyanobacterial diversity in lichen symbioses. MycoKeys. 2013.

66. Crits-Christoph A, Diamond S, Butterfield CN, Thomas BC, Banfield JF. Novel soil bacteria possess diverse genes for secondary metabolite biosynthesis. Nature. 2018;558(7710):440.

67. Chevrette MG, Carlson CM, Ortega HE, Thomas C, Ananiev GE, Barns KJ, et al. The antimicrobial potential of Streptomyces from insect microbiomes. Nat Commun. 2019;10(1):516.

68. Cragg GM, Newman DJ. Natural Products: a Continuing Source of Novel. Biochim Biophys Acta. 2013;1830(6):3670–95.

69. Payne DJ. Desperately seeking new antibiotics. Science. 2008;321(5896):1644–5.

70. Davies J, Wang H, Taylor T, Warabi K, Huang X-H, Andersen RJ. Uncialamycin, a new enediyne antibiotic. Organic letters. 2005;7(23):5233–6.

71. Parrot D, Antony-Babu S, Intertaglia L, Grube M, Tomasi S, Suzuki MT. Littoral lichens as a novel source of potentially bioactive Actinobacteria. Scientific Reports. 2015;5(1):15839-.

72. Barka EA, Vatsa P, Sanchez L, Gaveau-Vaillant N, Jacquard C, Klenk H-P, et al. Taxonomy, physiology, and natural products of Actinobacteria. Microbiology and Molecular Biology Molecular. 2016;80(1):1–43.

73. Liu C, Jiang Y, Wang X, Chen D, Chen X, Wang L, et al. Diversity, Antimicrobial Activity, and Biosynthetic Potential of Cultivable Actinomycetes Associated with Lichen Symbiosis. Microbial Ecology. 2017;74(3):570–84.

74. Selbmann L, Zucconi L, Ruisi S, Grube M, Cardinale M, Onofri S. Culturable bacteria associated with Antarctic lichens: Affiliation and psychrotolerance. Polar Biology. 2010;33(1):71–83.

75. Bolger AM, Lohse M, Usadel B. Trimmomatic: a flexible trimmer for Illumina sequence data. Bioinformatics. 2014;30(15):2114–20.

76. Schloss PD, Westcott SL, Ryabin T, Hall JR, Hartmann M, Hollister EB, et al. Introducing mothur: open-source, platform-independent, community-supported software for describing and comparing microbial communities. Appl Environ Microbiol. 2009;75(23):7537–41.

77. Quast C, Pruesse E, Yilmaz P, Gerken J, Schweer T, Yarza P, et al. The SILVA ribosomal RNA gene database project: improved data processing and web-based tools. Nucleic acids research. 2012;41(D1):D590–D6.

78. McDonald D, Price MN, Goodrich J, Nawrocki EP, DeSantis TZ, Probst A, et al. An improved Greengenes taxonomy with explicit ranks for ecological and evolutionary analyses of bacteria and archaea. The ISME journal. 2012;6(3):610.

79. Gloor G. ALDEx2: ANOVA-Like Differential Expression tool for compositional data. ALDEX manual modular. 2015;20.

80. Shannon CE. The mathematical theory of communication. 1963. MD Comput. 1997;14(4):306–17.

81. Simpson EH. Measurement of Diversity. Nature. 1949;163(4148):688-.

82. McMurdie PJ, Holmes S. phyloseq: an R package for reproducible interactive analysis and graphics of microbiome census data. PloS one. 2013;8(4):e61217.

83. Gloor G, Wong RG, Fernandes A, Albert A, Links M, Gloor MG, et al. Package ‘ALDEx2’. 2014.

84. Urbaniak C, McMillan A, Angelini M, Gloor GB, Sumarah M, Burton JP, et al. Effect of chemotherapy on the microbiota and metabolome of human milk, a case report. Microbiome. 2014;2(1):24.

85. Freitas AC, Bocking A, Hill JE, Money DM. Increased richness and diversity of the vaginal microbiota and spontaneous preterm birth. Microbiome. 2018;6(1):117.

86. Katoh K, Rozewicki J, Yamada KD. MAFFT online service: multiple sequence alignment, interactive sequence choice and visualization. Briefings in bioinformatics. 2017.

87. Katoh K, Misawa K, Kuma Ki, Miyata T. MAFFT: a novel method for rapid multiple sequence alignment based on fast Fourier transform. Nucleic acids research. 2002;30(14):3059–66.

88. Sokal RR. A statistical method for evaluating systematic relationship.. University of Kansas science bulletin; 1958. p. 1409–38.

89. González I, Ayuso-Sacido A, Anderson A, Genilloud O. Actinomycetes isolated from lichens: evaluation of their diversity and detection of biosynthetic gene sequences. FEMS microbiology ecology. 2005;54(3):401–15.

90. Wang D-s, Xue Q-h, Ma Y-y, Wei X-l, Chen J, He F. Oligotrophy is helpful for the isolation of bioactive actinomycetes. Indian journal of microbiology. 2014;54(2):178–84.

91. Mao DP, Zhou Q, Chen CY, Quan ZX. Coverage evaluation of universal bacterial primers using the metagenomic datasets. BMC Microbiol. 2012;12:66.

92. Bellemain E, Carlsen T, Brochmann C, Coissac E, Taberlet P, Kauserud H. ITS as an environmental DNA barcode for fungi: an in silico approach reveals potential PCR biases. BMC microbiology. 2010;10(1):189.

93. Hockett KL, Baltrus DA. Use of the Soft-agar Overlay Technique to Screen for Bacterially Produced Inhibitory Compounds. J Vis Exp. 2017(119).

94. Saffari N, Salmanzadeh-Ahrabi S, Abdi-Ali A, Rezaei-Hemami M. rA comparison of antibiotic disks from different sources on Quicolor and Mueller-Hinton agar media in evaluation of antibacterial susceptibility testing. Iranian journal of microbiology. 2016;8(5):307.

