## Supplemental Table 1 for "The microbiomes of seven lichen genera reveal host specificity, a reduced core community and potential as source of antimicrobials"

| Sample_code | Location | GPS | Lichen_Genus | Sustrate |
|---|---|---|---|---|
| Cla_B_Nev | Nevados | N 4.860833° W 75.361944° | *Cladonia* | TER |
| Cla_C_Nev | Nevados | N 4.909528° W 75.353694° | *Cladonia* | TER |
| Cla_D_Nev | Nevados | N 4.933056° W 75.355000° | *Cladonia* | COR |
| Cla_E_Nev | Nevados | N 4.968523° W 75.352512° | *Cladonia* | TER |
| Cla_F_Nev | Nevados | N 4.970833° W 75.351944° | *Cladonia* | TER |
| Co_B_Nev | Nevados | N 4.860833° W 75.361944° | *Cora* | TER |
| Co_F_Nev | Nevados | N 4.970833° W 75.351944° | *Cora* | TER |
| Hyp_C_Nev | Nevados | N 4.909528° W 75.353694° | *Hypotrachyna* | COR |
| Hyp_E_Nev | Nevados | N 4.968523° W 75.352512° | *Hypotrachyna* | COR |
| Hyp_F_Nev | Nevados | N 4.970833° W 75.351944° | *Hypotrachyna* | COR |
| Ptg_F_Nev | Nevados | N 4.970833° W 75.351944° | *Peltigera* | TER |
| Stc_A_Nev | Nevados | N 4.852778° W 75.364722° | *Stereocaulon* | TER |
| Stc_B_Nev | Nevados | N 4.860833° W 75.361944° | *Stereocaulon* | TER |
| Stc_D_Nev | Nevados | N 4.933056° W 75.355000° | *Stereocaulon* | TER |
| Stc_F_Nev | Nevados | N 4.970833° W 75.351944° | *Stereocaulon* | TER |
| Stt_C_Nev | Nevados | N 4.909528° W 75.353694° | *Sticta* | COR |
| Stt_D_Nev | Nevados | N 4.933056° W 75.355000° | *Sticta* | TER |
| Us_A_Nev | Nevados | N 4.852778° W 75.364722° | *Usnea* | COR |
| Us_C_Nev | Nevados | N 4.909528° W 75.353694° | *Usnea* | COR |
| Us_D_Nev | Nevados | N 4.933056° W 75.355000° | *Usnea* | COR |
| Us_E_Nev | Nevados | N 4.968523° W 75.352512° | *Usnea* | COR |
| Cla_A_Chi | Chingaza | N 4.71470° W 73.82208° | *Cladonia* | SAX |
| Cla_K1_Chi | Chingaza | N 4.67727° W 73.78642° | *Cladonia* | TER |
| Cla_K2_Chi | Chingaza | N 4.67727° W 73.78642° | *Cladonia* | TER |
| Cla_L_Chi | Chingaza | N 4.67766° W 73.78671° | *Cladonia* | TER |
| Co_A_Chi | Chingaza | N 4.71470° W 73.82208° | *Cora* | SAX |
| Co_H1_Chi | Chingaza | N 4.62904° W 73.67063° | *Cora* | COR |
| Co_H2_Chi | Chingaza | N 4.62904° W 73.67063° | *Cora* | COR |
| Co_I_Chi | Chingaza | N 4.62377° W 73.72462° | *Cora* | TER |
| Co_L_Chi | Chingaza | N 4.67766° W 73.78671° | *Cora* | TER |
| Hyp_B_Chi | Chingaza | N 4.62480° W 73.72694° | *Hypotrachyna* | COR |
| Hyp_H_Chi | Chingaza | N 4.62904° W 73.67063° | *Hypotrachyna* | TER |
| Hyp_M_Chi | Chingaza | N 4.68166° W 73.78699° | *Hypotrachyna* | SAX |
| Hyp_N_Chi | Chingaza | N 4.68298° W 73.78704° | *Hypotrachyna* | SAX |
| Ptg_C_Chi | Chingaza | N 4.62426° W 73.72611° | *Peltigera* | COR |
| Ptg_N_Chi | Chingaza | N 4.68298° W 73.78704° | *Peltigera* | SAX |
| Stc_A_Chi | Chingaza | N 4.71470° W 73.82208° | *Stereocaulon* | SAX |
| Stc_J_Chi | Chingaza | N 4.68306° W 73.78687° | *Stereocaulon* | SAX |
| Stc_L_Chi | Chingaza | N 4.67766° W 73.78671° | *Stereocaulon* | TER |
| Stt_A_Chi | Chingaza | N 4.71470° W 73.82208° | *Sticta* | COR |
| Stt_B_Chi | Chingaza | N 4.62480° W 73.72694° | *Sticta* | COR |
| Stt_C_Chi | Chingaza | N 4.62426° W 73.72611° | *Sticta* | COR |
| Stt_E_Chi | Chingaza | N 4.62349° W 73.72566° | *Sticta* | COR |
| Stt_F_Chi | Chingaza | N 4.623400° W 73.725590° | *Sticta* | COR |
| Us_E_Chi | Chingaza | N 4.62349° W 73.72566° | *Usnea* | COR |
| Us_G_Chi | Chingaza | N 4.62335° W 73.72551° | *Usnea* | COR |
| Us_K_Chi | Chingaza | N 4.67727° W 73.78642° | *Usnea* | COR |
