## Supplemental Table 2 for "The microbiomes of seven lichen genera reveal host specificity, a reduced core community and potential as source of antimicrobials"

| Samples | Lichen | Observed | Shannon | Simpson |
|---|---|---|---|---|
| Cla_A_Chi | Cladonia | 769 | 4.13345805 | 0.94352985 |
| Cla_B_Nev | Cladonia | 324 | 1.83803213 | 0.71642585 |
| Cla_C_Nev | Cladonia | 208 | 1.68723544 | 0.71813521 |
| Cla_D_Nev | Cladonia | 260 | 2.23916555 | 0.76769596 |
| Cla_E_Nev | Cladonia | 385 | 2.65079485 | 0.83552612 |
| Cla_F_Nev | Cladonia | 298 | 1.61505157 | 0.59386502 |
| Cla_K1_Chi | Cladonia | 567 | 3.61279749 | 0.92669056 |
| Cla_K2_Chi | Cladonia | 868 | 4.29626725 | 0.96763313 |
| Cla_L_Chi | Cladonia | 1106 | 4.58981752 | 0.96913622 |
| Co_A_Chi | Cora | 996 | 3.47136917 | 0.78522872 |
| Co_B_Nev | Cora | 201 | 1.68133916 | 0.68316316 |
| Co_F_Nev | Cora | 278 | 1.63636832 | 0.64028134 |
| Co_H1_Chi | Cora | 1312 | 4.57092575 | 0.93849366 |
| Co_H2_Chi | Cora | 767 | 3.29613314 | 0.83855433 |
| Co_I_Chi | Cora | 210 | 1.83993806 | 0.6572326 |
| Co_L_Chi | Cora | 758 | 3.22255803 | 0.8290168 |
| Hyp_B_Chi | Hypotrachyna | 819 | 4.62616387 | 0.97363313 |
| Hyp_C_Nev | Hypotrachyna | 619 | 2.29316784 | 0.58821926 |
| Hyp_E_Nev | Hypotrachyna | 427 | 3.74761329 | 0.9548576 |
| Hyp_F_Nev | Hypotrachyna | 782 | 4.73816764 | 0.97990882 |
| Hyp_H_Chi | Hypotrachyna | 1924 | 5.61251287 | 0.9854916 |
| Hyp_M_Chi | Hypotrachyna | 704 | 4.01967358 | 0.94366012 |
| Hyp_N_Chi | Hypotrachyna | 1003 | 4.20067332 | 0.89489018 |
| Ptg_C_Chi | Peltigera | 1955 | 4.68707721 | 0.88646336 |
| Ptg_F_Nev | Peltigera | 433 | 2.02338715 | 0.63823252 |
| Ptg_N_Chi | Peltigera | 639 | 2.31384759 | 0.62289145 |
| Stc_A_Chi | Stereocaulon | 661 | 3.43656293 | 0.87841948 |
| Stc_A_Nev | Stereocaulon | 343 | 1.94275398 | 0.57475456 |
| Stc_B_Nev | Stereocaulon | 843 | 3.58251167 | 0.85479688 |
| Stc_D_Nev | Stereocaulon | 352 | 1.87980636 | 0.62319748 |
| Stc_F_Nev | Stereocaulon | 220 | 1.75184541 | 0.7101174 |
| Stc_J_Chi | Stereocaulon | 738 | 3.75418307 | 0.90163427 |
| Stc_L_Chi | Stereocaulon | 882 | 4.38191555 | 0.95311243 |
| Stt_A_Chi | Sticta | 1611 | 4.45935331 | 0.86047591 |
| Stt_B_Chi | Sticta | 1078 | 3.89970208 | 0.82525909 |
| Stt_C_Chi | Sticta | 1418 | 5.24448878 | 0.98379839 |
| Stt_C_Nev | Sticta | 504 | 2.33674598 | 0.71590542 |
| Stt_D_Nev | Sticta | 476 | 1.7477335 | 0.56932314 |
| Stt_E_Chi | Sticta | 1197 | 5.10014433 | 0.98241538 |
| Stt_F_Chi | Sticta | 1633 | 5.39393032 | 0.98609535 |
| Us_A_Nev | Usnea | 100 | 0.56393615 | 0.20865696 |
| Us_C_Nev | Usnea | 189 | 0.89319927 | 0.26074171 |
| Us_D_Nev | Usnea | 425 | 2.23491125 | 0.77018313 |
| Us_E_Chi | Usnea | 763 | 3.19272753 | 0.73691332 |
| Us_E_Nev | Usnea | 177 | 2.20428949 | 0.69595366 |
| Us_G_Chi | Usnea | 318 | 2.68682305 | 0.87396244 |
| Us_K_Chi | Usnea | 185 | 2.06754498 | 0.77856529 |

| Alpha | 0.05 | | | | |
|---|---|---|---|---|---|
| **Tukey's multiple comparisons test** | **Mean Diff.** | **95.00% CI of diff.** | **Significant?** | **Summary** | **Adjusted P Value** |
| Cladonia vs. Cora | 0.05909 | -0.1999 to 0.3181 | No | ns | 0.9913 |
| Cladonia vs. Hypotrachyna | -0.07644 | -0.3354 to 0.1825 | No | ns | 0.9678 |
| Cladonia vs. Peltigera | 0.1107 | -0.2319 to 0.4532 | No | ns | 0.9505 |
| Cladonia vs. Stereocaulon | 0.04137 | -0.2176 to 0.3003 | No | ns | 0.9988 |
| Cladonia vs. Sticta | -0.01967 | -0.2786 to 0.2393 | No | ns | >0.9999 |
| Cladonia vs. Usnea | 0.2087 | -0.05032 to 0.4676 | No | ns | 0.1866 |
| Cora vs. Hypotrachyna | -0.1355 | -0.4102 to 0.1392 | No | ns | 0.7249 |
| Cora vs. Peltigera | 0.05156 | -0.3031 to 0.4062 | No | ns | 0.9993 |
| Cora vs. Stereocaulon | -0.01772 | -0.2924 to 0.2570 | No | ns | >0.9999 |
| Cora vs. Sticta | -0.07876 | -0.3534 to 0.1959 | No | ns | 0.9721 |
| Cora vs. Usnea | 0.1496 | -0.1251 to 0.4243 | No | ns | 0.6266 |
| Hypotrachyna vs. Peltigera | 0.1871 | -0.1675 to 0.5417 | No | ns | 0.6599 |
| Hypotrachyna vs. Stereocaulon | 0.1178 | -0.1569 to 0.3925 | No | ns | 0.8334 |
| Hypotrachyna vs. Sticta | 0.05677 | -0.2179 to 0.3315 | No | ns | 0.9949 |
| Hypotrachyna vs. Usnea | 0.2851 | 0.01041 to 0.5598 | Yes | * | 0.0375 |
| Peltigera vs. Stereocaulon | -0.06929 | -0.4239 to 0.2853 | No | ns | 0.9962 |
| Peltigera vs. Sticta | -0.1303 | -0.4849 to 0.2243 | No | ns | 0.9115 |
| Peltigera vs. Usnea | 0.09801 | -0.2566 to 0.4526 | No | ns | 0.9767 |
| Stereocaulon vs. Sticta | -0.06103 | -0.3357 to 0.2137 | No | ns | 0.9925 |
| Stereocaulon vs. Usnea | 0.1673 | -0.1074 to 0.4420 | No | ns | 0.4985 |
| Sticta vs. Usnea | 0.2283 | -0.04636 to 0.5030 | No | ns | 0.1597 |

| Alpha | 0.05 | | | | |
|---|---|---|---|---|---|
| **Tamhane's T2 multiple comparisons test** | **Mean Diff.** | **95.00% CI of diff.** | **Significant?** | **Summary** | **djusted P Value** |
| Cladonia vs. Cora | 0.1456 | -2.022 to 2.313 | No | ns | >0.9999 |
| Cladonia vs. Hypotrachyna | -1.214 | -3.277 to 0.8481 | No | ns | 0.6389 |
| Cladonia vs. Peltigera | -0.04559 | -9.108 to 9.017 | No | ns | >0.9999 |
| Cladonia vs. Stereocaulon | 0.001145 | -2.113 to 2.115 | No | ns | >0.9999 |
| Cladonia vs. Sticta | -1.064 | -3.689 to 1.562 | No | ns | 0.9631 |
| Cladonia vs. Usnea | 0.9849 | -0.9874 to 2.957 | No | ns | 0.8517 |
| Cora vs. Hypotrachyna | -1.36 | -3.559 to 0.8396 | No | ns | 0.5348 |
| Cora vs. Peltigera | -0.1912 | -8.915 to 8.533 | No | ns | >0.9999 |
| Cora vs. Stereocaulon | -0.1444 | -2.387 to 2.098 | No | ns | >0.9999 |
| Cora vs. Sticta | -1.209 | -3.909 to 1.491 | No | ns | 0.9118 |
| Cora vs. Usnea | 0.8393 | -1.287 to 2.966 | No | ns | 0.9711 |
| Hypotrachyna vs. Peltigera | 1.169 | -8.167 to 10.50 | No | ns | 0.9994 |
| Hypotrachyna vs. Stereocaulon | 1.215 | -0.9323 to 3.363 | No | ns | 0.6705 |
| Hypotrachyna vs. Sticta | 0.1508 | -2.500 to 2.801 | No | ns | >0.9999 |
| Hypotrachyna vs. Usnea | 2.199 | 0.1862 to 4.212 | Yes | * | 0.0268 |
| Peltigera vs. Stereocaulon | 0.04674 | -8.970 to 9.064 | No | ns | >0.9999 |
| Peltigera vs. Sticta | -1.018 | -8.161 to 6.126 | No | ns | >0.9999 |
| Peltigera vs. Usnea | 1.03 | -8.971 to 11.03 | No | ns | 0.9999 |
| Stereocaulon vs. Sticta | -1.065 | -3.738 to 1.609 | No | ns | 0.9655 |
| Stereocaulon vs. Usnea | 0.9837 | -1.085 to 3.053 | No | ns | 0.8727 |
| Sticta vs. Usnea | 2.048 | -0.5668 to 4.664 | No | ns | 0.1984 |

| Alpha | 0.05 | | | | |
|---|---|---|---|---|---|
| **Tukey's multiple comparisons t** | **Mean Diff.** | **.00% CI of di** | **Significant?** | **Summary** | **djusted P Value** |
| Cladonia vs. Cora | -114.3 | 760.7 to 532. | No | ns | 0.9978 |
| Cladonia vs. Hypotrachyna | -365.2 | 1012 to 281. | No | ns | 0.5859 |
| Cladonia vs. Peltigera | -477.3 | 1332 to 377. | No | ns | 0.5993 |
| Cladonia vs. Stereocaulon | -45.33 | 691.7 to 601. | No | ns | >0.9999 |
| Cladonia vs. Sticta | -599.3 | 1246 to 47.0 | No | ns | 0.0847 |
| Cladonia vs. Usnea | 223.5 | 422.8 to 869. | No | ns | 0.9323 |
| Cora vs. Hypotrachyna | -250.9 | 936.4 to 434. | No | ns | 0.9131 |
| Cora vs. Peltigera | -363 | 1248 to 522. | No | ns | 0.8601 |
| Cora vs. Stereocaulon | 69 | 616.6 to 754. | No | ns | >0.9999 |
| Cora vs. Sticta | -485 | 1171 to 200. | No | ns | 0.3207 |
| Cora vs. Usnea | 337.9 | 347.7 to 102 | No | ns | 0.726 |
| Hypotrachyna vs. Peltigera | -112.1 | 997.2 to 772. | No | ns | 0.9997 |
| Hypotrachyna vs. Stereocaulon | 319.9 | 365.7 to 100 | No | ns | 0.7728 |
| Hypotrachyna vs. Sticta | -234.1 | 919.7 to 451. | No | ns | 0.936 |
| Hypotrachyna vs. Usnea | 588.7 | 96.87 to 127 | No | ns | 0.1342 |
| Peltigera vs. Stereocaulon | 432 | 453.1 to 131 | No | ns | 0.7346 |
| Peltigera vs. Sticta | -122 | 1007 to 763. | No | ns | 0.9995 |
| Peltigera vs. Usnea | 700.9 | 184.2 to 158 | No | ns | 0.2025 |
| Stereocaulon vs. Sticta | -554 | 1240 to 131. | No | ns | 0.184 |
| Stereocaulon vs. Usnea | 268.9 | 416.7 to 954. | No | ns | 0.8834 |
| Sticta vs. Usnea | 822.9 | 137.3 to 150 | Yes | ** | 0.01 |
