## Supplemental Table 3 for "The microbiomes of seven lichen genera reveal host specificity, a reduced core community and potential as source of antimicrobials"

| GenBank-ID | Phylum | Order | Lichen | Continent | Country | Biome |
|---|---|---|---|---|---|---|
| MG996731_1 | Proteobacteria | Rhizobiales | Yes | Europe | Russia | NA |
| GU300236_1 | Proteobacteria | Rhizobiales | No | America | Canada | Forest |
| JX967285_1 | NA | NA | Yes | Europe | Norway | Arctic |
| HQ333277_1 | NA | NA | No | Asia | China | Glacier |
| HM240934_1 | NA | NA | No | NA | NA | Desert |
| LN614695_1 | Proteobacteria | Rhizobiales | No | America | USA | Glacier |
| JX967294_1 | Proteobacteria | Rhizobiales | Yes | Europe | Norway | Arctic |
| AM940729_1 | Proteobacteria | Rhodospirillales | No | Europe | Norway | Arctic |
| JN023390_1 | NA | NA | Yes | America | Mexico | Grassland |
| LN929548_1 | NA | NA | No | America | Chile | Desert |
| JX098367_1 | NA | NA | No | America | Chile | Desert |
| JN023294_1 | NA | NA | Yes | America | Mexico | Grassland |
| FR749823_1 | Acidobacteria | NA | No | Antarctica | NA | Antarctic |
| KT752896_1 | Acidobacteria | NA | No | Europe | Norway | Glacier |
| EU083407_1 | NA | NA | No | Europe | Switzerland | Glacier |
| JQ381812_2 | NA | NA | No | America | USA | NA |
| NR_118023_1 | Acidobacteria | Acidobacteriales | No | Europe | Finland | Tundra |
| HQ595216_1 | Acidobacteria | | No | Europe | Norway | Arctic |
| JQ480554_1 | NA | NA | No | Europe | Switzerland | Glacier |
| EF220623_1 | Proteobacteria | NA | No | America | Falkland Islands | Antarctic |
| KU219090_1 | Cyanobacteria | NA | No | America | Brazil | Forest |
| EU861940_1 | NA | NA | No | America | USA | Tundra |
| EF221408_1 | Proteobacteria | NA | No | Antarctica | NA | Antarctic |
| HF546516_1 | Acidobacteria | NA | Yes | Europe | Italy | Glacier |
| HF546512_1 | Proteobacteria | Rhodospirillales | Yes | Europe | Italy | Glacier |
| HF546509_1 | Cyanobacteria | NA | Yes | Europe | Italy | Glacier |
| FJ475466_1 | Proteobacteria | Rhodospirillales | No | Europe | Sweden | Forest |
| KX467876_1 | Proteobacteria | Rhizobiales | No | Europe | Norway | Arctic |
| MH530970_1 | NA | NA | No | Australia | New Zeland | Forest |
| JF907343_1 | NA | NA | No | America | Argentina | Antarctic |
| KU219013_1 | Cyanobacteria | NA | No | America | Brazil | Forest |
| JX967352_1 | NA | NA | Yes | Europe | Norway | Arctic |
| MF423481_1 | Cyanobacteria | Nostocales | Yes | America | USA | Forest |
| AF428513_1 | Cyanobacteria | NA | Yes | America | USA | Desert |
| Otu00008 | Proteobacteria | Rhizobiales | Yes | America | Colombia | Paramo |
| Otu00012 | Proteobacteria | Rhizobiales | Yes | America | Colombia | Paramo |
| Otu00015 | Proteobacteria | Rhodospirillales | Yes | America | Colombia | Paramo |
| Otu00016 | Proteobacteria | Sphingomonadales | Yes | America | Colombia | Paramo |
| Otu00017 | Acidobacteria | Acidobacteriales | Yes | America | Colombia | Paramo |
| Otu00018 | Acidobacteria | Acidobacteriales | Yes | America | Colombia | Paramo |
| Otu00019 | Proteobacteria | Rhizobiales | Yes | America | Colombia | Paramo |
| Otu00026 | Proteobacteria | Rhizobiales | Yes | America | Colombia | Paramo |
| Otu00032 | Proteobacteria | Rhodospirillales | Yes | America | Colombia | Paramo |
| Otu00036 | Cyanobacteria | NA | Yes | America | Colombia | Paramo |
| Otu00037 | Proteobacteria | Rhodospirillales | Yes | America | Colombia | Paramo |
| Otu00041 | Acidobacteria | Acidobacteriales | Yes | America | Colombia | Paramo |
| Otu00042 | Proteobacteria | Rhodospirillales | Yes | America | Colombia | Paramo |
| Otu00047 | Proteobacteria | Rhodospirillales | Yes | America | Colombia | Paramo |
| Otu00062 | Proteobacteria | Rhizobiales | Yes | America | Colombia | Paramo |
| Otu00112 | Acidobacteria | Acidobacteriales | Yes | America | Colombia | Paramo |
