## Supplemental Table 4 for "The microbiomes of seven lichen genera reveal host specificity, a reduced core community and potential as source of antimicrobials"

| Lichen | Code | Medium | Growth time | Macroscopic Description | | | | Bioactive | Against | PCR 16S rRNA (27F-1492R) | Microorganism |
|---|---|---|---|---|---|---|---|---|---|---|---|
| | | | | Color | Surface | Shape | Edge | | | | |
| | PN.F6.1 | ISP2 | 24h | Orange | Smooth | Irregular | Irregular | Yes | *C. albicans* | Positive | Bacteria |
| *Bacidia* | PN.F6.2 | ISP2 | 3 Days | Beige | Rough | Irregular | Irregular | No | NA | Positive | Bacteria |
| | PN.F6.3 | DIFCO | 24h | White | Smooth | Irregular | Irregular | Yes | *S. aureus, A. baumannii* | Positive | Bacteria |
| | PN.F6.4 | DIFCO | >5 Days | White | Smooth | Irregular | Irregular | No | NA | Positive | Bacteria |
| | PN.C2 | ISP2 | >5 Days | White | Rough | Punctiform | Regular | No | NA | Positive | Bacteria |
| | PN.C2.0 | ISP2 | >5 Days | White | Rough | Circular | Regular | No | NA | Positive | Bacteria |
| | PN.C2.2 | DIFCO | >5 Days | Beige | Rough | Irregular | Irregular | No | NA | Positive | Bacteria |
| | PN.C2.3 | ISP2 | >5 Days | Pink | Smooth | Irregular | Irregular | No | NA | Positive | Bacteria |
| *Cladonia* | PN.C2.4 | GAUSE | >5 Days | White | Smooth | Circular | Regular | No | NA | Positive | Bacteria |
| | PN.C2.5 | ISP2 | >5 Days | Beige | Rough | Circular | Regular | No | NA | Positive | Bacteria |
| | PN.D1.1 | ISP2 | >5 Days | Green | Smooth | Punctiform | Irregular | No | NA | Positive | Bacteria |
| | PN.D1.2 | ISP2 | >5 Days | Brown | Smooth | Irregular | Irregular | No | NA | Positive | Bacteria |
| | PN.E1.1 | DIFCO | >5 Days | Yellow | Smooth | Circular | Regular | No | NA | Positive | Bacteria |
| | PN.E1.2 | ISP2 | >5 Days | Green | Smooth | Irregular | Regular | No | NA | Positive | Bacteria |
| | PN.F1.1 | ISP2 | 72h | Yellow | Rough | Irregular | Irregular | No | NA | Positive | Bacteria |
| | PN.F1.3 | DIFCO | >5 Days | Pink | Rough | Punctiform | Regular | No | NA | Positive | Bacteria |
| *Cora* | PN.F1.4 | DIFCO | >5 Days | Pink | Rough | Irregular | Irregular | Yes | *A.baumannii* | Positive | Bacteria |
| | PN.F1.6 | DIFCO | >5 Days | No color | Rough | Punctiform | Regular | No | NA | Positive | Bacteria |
| | PN.F1.9 | GAUSE | >5 Days | White | Smooth | Irregular | Irregular | No | NA | Positive | Bacteria |
| | PN.C6.2 | DIFCO | 48h | White | Smooth | Irregular | Regular | No | NA | Positive | Bacteria |
| *Hypotrachyna* | PN.F3.1 | ISP2 | >5 Days | Beige | Smooth | Irregular | Irregular | No | NA | Positive | Bacteria |
| | PN.F3.2 | DIFCO | >5 Days | Pink | Rough | Punctiform | Regular | No | NA | Positive | Bacteria |
| | PN.F4.1 | ISP2 | >5 Days | Orange | Rough | Punctiform | Irregular | No | NA | Positive | Bacteria |
| | PN.F4.10 | DIFCO | >5 Days | White | Smooth | Punctiform | Regular | No | NA | Positive | Bacteria |
| *Peltigera* | PN.F4.3 | ISP2 | >5 Days | Brown | Smooth | Irregular | Irregular | Yes | *S. aureus, C. albicans, P. aeruginosa* | Positive | Bacteria |
| | PN.F4.7 | DIFCO | >5 Days | Beige | Smooth | Irregular | Irregular | Yes | *S. aureus, P. aeruginosa* | Positive | Bacteria |
| | PN.F4.9 | DIFCO | >5 Days | Orange | Smooth | Irregular | Irregular | No | NA | Positive | Bacteria |
| *Psoroma* | PN.C4.3 | ISP2 | >5 Days | White | Smooth | Punctiform | Irregular | Yes | *K. pneumoniae* | Positive | Bacteria |
| | PN.C4.4 | DIFCO | >5 Days | Pink | Rough | Punctiform | Irregular | No | NA | Positive | Bacteria |
| | PN.A3.1 | DIFCO | >5 Days | White | Rough | Irregular | Irregular | No | NA | Positive | Bacteria |
| | PN.A3.3 | ISP2 | >5 Days | Cream | Rough | Irregular | Irregular | No | NA | Positive | Bacteria |
| | PN.A3.4 | DIFCO | >5 Days | Yellow | Smooth | Punctiform | Regular | No | NA | Positive | Bacteria |
| *Stereocaulon* | PN.A3.5 | ISP2 | >5 Days | Beige | Smooth | Punctiform | Regular | No | NA | Positive | Bacteria |
| | PN.B1.3 | ISP2 | >5 Days | Pink | Smooth | Punctiform | Irregular | No | NA | Positive | Bacteria |
| | PN.B1.4 | ISP2 | >5 Days | Yellow | Rough | Circular | Irregular | No | NA | Positive | Bacteria |
| | PN.D2.2 | DIFCO | >5 Days | White | Smooth | Irregular | Irregular | No | NA | Positive | Bacteria |
| | PN.F5.2 | DIFCO | >5 Days | White | Smooth | Punctiform | Irregular | Yes | *C.albicans, A. baumannii* | Positive | Bacteria |
| | PN.C3.1 | ISP2 | 72h | Yellow | Rough | Punctiform | Irregular | Yes | *S. enterica, E.coli, P. aeuriginosa* | Positive | Bacteria |
| | PN.C3.2 | DIFCO | >5 Days | Red | Smooth | Punctiform | Regular | Yes | *C. albicans* | Positive | Bacteria |
| *Sticta* | PN.C3.3 | ISP2 | 48h | Beige | Rough | Circular | Regular | No | NA | Positive | Bacteria |
| | PN.D3.1 | DIFCO | >5 Days | Red | Rough | Irregular | Regular | No | NA | Positive | Bacteria |
| | PN.D3.3 | ISP2 | >5 Days | Yellow | Smooth | Irregular | Irregular | No | NA | Positive | Bacteria |
| | PN.D3.4 | DIFCO | >5 Days | Yellow | Smooth | Irregular | Irregular | No | NA | Positive | Bacteria |
| | PN.A2.1 | DIFCO | >5 Days | Pink | Smooth | Punctiform | Irregular | No | NA | Positive | Bacteria |
| *Usnea* | PN.D4.1 | ISP2 | >5 Days | Beige | Rough | Circular | Regular | No | NA | Positive | Bacteria |
| | PN.D4.2 | ISP2 | >5 Days | White | Rough | Irregular | Regular | No | NA | Positive | Bacteria |

ISP2: Streptomyces Project Medium-2
DIFCO: Actinomycete Isolation Agar
GAUSE: Gause Synthetic Agar

Description code name: PN corresponds to Nevados Park; The following letter represents the sublocation point with the number of lichen thalli; Following number represents the numeration for the isolate

| Lichen | Code | Medium | Growth time | Macroscopic Description | | | | Bioactive | Against | PCR 16S rRNA (27F-1492R) | PCR ITS (ITS4-IT5) | Microorganism |
|---|---|---|---|---|---|---|---|---|---|---|---|---|
| | | | | Color | Surface | Shape | Edge | | | | | |
| *Cladonia* | PC.L1.2 | DIFCO | | NA | NA | NA | NA | No | NA | Negative | Positive | Fungi |
| | PC.L1.3 | DIFCO | | NA | NA | NA | NA | No | NA | Negative | Positive | Fungi |
| | PC.A4.4 | ISP2 | 4 Days | White | Rough | Irregular | Irregular | Yes | *S. aureus, A. baumannii* | Positive | | Bacteria |
| | PC.A4.5 | GAUSE | | White | Rough | Circular | Irregular | Yes | *S. aureus, A. baumannii* | Positive | | Bacteria |
| *Yoshimuriella* | PC.C1.1 | ISP2 | | White | Rough | Circular | Irregular | Yes | *K. pneumoniae* | Positive | | Bacteria |
| | PC.C1.7 | GAUSE-O | >5 Days | White | Rough | Circular | Regular | No | NA | Positive | | Bacteria |
| | PC.C1.8 | GAUSE | | NA | NA | NA | NA | No | NA | Positive | | Bacteria |
| | PC.C1.9 | GAUSE | | NA | NA | NA | NA | No | NA | Negative | Positive | Fungi |
| | PC.D1.5 | ISP2 | 4 Days | Yellow | Smooth | Punctiform | Regular | No | NA | Positive | | Bacteria |
| | PC.D1.6 | GAUSE-O | 4 Days | White | Rough | Circular | Irregular | No | NA | Positive | | Bacteria |
| | PC.D1.7 | GAUSE | | Cream | Rough | Circular | Regular | No | NA | Positive | | Bacteria |
| | PC.D1.8 | ISP2 | | NA | NA | NA | NA | Yes | *S. aureus, A. baumannii* | Positive | | Bacteria |
| | PC.D1.10 | GAUSE-O | | White | Rough | Circular | Regular | No | NA | Positive | | Bacteria |
| | PC.D1.11 | DIFCO | | Brown | Rough | Circular | Irregular | No | NA | Positive | | Bacteria |
| | PC.D1.12 | ISP2 | | White | Rough | Circular | Irregular | No | NA | Positive | | Bacteria |
| *Sticta* | PC.E2.3 | ISP2 | >5 Days | White | Rough | Irregular | Irregular | No | NA | Positive | | Bacteria |
| | PC.E2.4 | ISP2 | >5 Days | White | Rough | Irregular | Irregular | NA | NA | Positive | | Bacteria |
| | PC.E2.5 | ISP2 | | White | Rough | Circular | Irregular | No | NA | Positive | | Bacteria |
| | PC.E2.6 | ISP2 | | NA | NA | NA | NA | No | NA | Positive | | Bacteria |
| | PC.E2.7 | DIFCO | | White | Rough | Circular | Irregular | Yes | *C. albicans* | Positive | | Bacteria |
| | PC.E2.8 | GAUSE | | White | Rough | Punctiform | Irregular | No | NA | Positive | | Bacteria |
| | PC.E2.9 | GAUSE | | White | Rough | Circular | Irregular | No | NA | Positive | | Bacteria |
| | PC.A5.7 | ISP2 | | Brown | Rough | Circular | Irregular | Yes | *S. aureus, A. baumannii* | Negative | Positive | Fungi |
| | PC.A5.8 | ISP2 | | NA | NA | NA | NA | Yes | *S. aureus, A. baumannii* | Positive | | Bacteria |
| *Lobariella** | PC.G1.1 | DIFCO | | NA | NA | NA | NA | No | NA | Negative | Positive | Fungi |
| | PC.G1.2 | DIFCO | | Black | Rough | Irregular | Irregular | No | NA | Negative | Positive | Fungi |
| | PC.G1.4 | DIFCO | | NA | NA | NA | NA | No | NA | Positive | | Bacteria |
| *Hypotrachyna* | PC.B1.1 | ISP2 | >5 Days | Cream | Smooth | Circular | Regular | No | NA | Positive | | Bacteria |
| | PC.B1.2 | DIFCO | | NA | NA | NA | NA | No | NA | Positive | | Bacteria |
| | PC.G2.1 | DIFCO | | White | Rough | Circular | Irregular | Yes | *A.baumannii* | Positive | | Bacteria |
| | PC.G2.2 | ISP2 | | Green | Smooth | Circular | Regular | No | NA | Positive | | Bacteria |
| | PC.G2.3 | ISP2 | | Cream | Smooth | Irregular | Irregular | No | NA | Negative | Positive | Fungi |
| | PC.H3.10 | ISP2 | 4 Days | Brown | Rough | Irregular | Irregular | No | NA | Positive | | Bacteria |
| | PC.H3.11 | ISP2 | 4 Days | Cream | Rough | Punctiform | Irregular | Yes | *S. aureus, A. baumannii, C. albicans* | NA | | Bacteria |
| | PC.H3.12 | ISP2 | | Yellow | Rough | Circular | Irregular | Yes | *S. aureus, A. baumannii* | Positive | | Bacteria |
| | PC.H3.13 | ISP2 | | Yellow | Smooth | Punctiform | Regular | Yes | *S. aureus, A. baumannii* | Positive | | Bacteria |
| | PC.H3.6 | DIFCO | 4 Days | Brown | Rough | Circular | Irregular | No | NA | Positive | | Bacteria |
| | PC.H3.7 | GAUSE | 4 Days | Brown | Rough | Irregular | Irregular | No | NA | Positive | | Bacteria |
| | PC.H3.8 | ISP2 | | Brown | Rough | Irregular | Irregular | No | NA | Positive | | Bacteria |
| | PC.H3.9 | ISP2 | 4 Days | White | Rough | Punctiform | Irregular | Yes | *C. albicans* | Positive | | Bacteria |
| | PC.H3.14 | DIFCO | | Yellow | Rough | Circular | Irregular | Yes | *A.baumannii* | Positive | | Bacteria |
| | PC.H3.15 | DIFCO | >5 Days | White | Rough | Circular | Irregular | No | NA | Positive | | Bacteria |
| | PC.H3.16 | ISP2 | | Yellow | Rough | Circular | Irregular | Yes | *S. aureus, A. baumannii, C. albicans* | Positive | | Bacteria |
| | PC.H3.17 | GAUSE | | Yellow | Rough | Punctiform | Irregular | Yes | *S. aureus, A. baumannii* | Positive | | Bacteria |
| | PC.H3.18 | DIFCO | | NA | NA | NA | NA | Yes | *S. aureus, A. baumannii* | Positive | | Bacteria |
| | PC.H3.19 | GAUSE-O | | NA | NA | NA | NA | Yes | *S. aureus, A. baumannii, C. albicans, F. oxysporum* | Positive | | Bacteria |
| | PC.N3.10 | DIFCO | 4 Days | Yellow | Smooth | Punctiform | Irregular | No | NA | Positive | | Bacteria |
| *Cora* | PC.H1.1 | ISP2 | >5 Days | Cream | Smooth | Irregular | Irregular | No | NA | Negative | Positive | Fungi |
| | PC.H2.1 | DIFCO | >5 Days | Yellow | Rough | Punctiform | Irregular | Yes | *S. aureus* | Positive | | Bacteria |
| | PC.H2.2 | ISP2 | | Orange | Smooth | Irregular | Irregular | Yes | *S. aureus, A. baumannii* | Negative | Positive | Fungi |
| | PC.H2.3 | ISP2 | | Cream | Smooth | Irregular | Irregular | No | NA | Positive | | Bacteria |
| | PC.I1.1 | DIFCO | | White | Smooth | Punctiform | Regular | Yes | *C. albicans* | Positive | | Bacteria |
| | PC.L2.1 | GAUSE-O | | NA | NA | NA | NA | Yes | *F. oxysporum* | Positive | | Bacteria |
| | PC.A3.3 | DIFCO | 4 Days | White | Rough | Circular | Irregular | No | NA | Positive | | Bacteria |
| | PC.A3.4 | ISP2 | | Cream | Smooth | Irregular | Irregular | No | NA | Positive | | Bacteria |
| *Stereocaulon* | PC.L3.2 | ISP2 | | White | Rough | Irregular | Irregular | No | NA | Positive | | Bacteria |
| | PC.L3.3 | ISP2 | | White | Rough | Punctiform | Irregular | No | NA | Positive | | Bacteria |
| | PC.L3.5 | ISP2 | | White | Rough | Punctiform | Irregular | No | NA | Positive | | Bacteria |
| | PC.L3.6 | ISP2 | | White | Rough | Punctiform | Irregular | No | NA | Positive | | Bacteria |
| | PC.L3.8 | DIFCO | | White | Rough | Circular | Regular | No | NA | Positive | | Bacteria |
| | PC.L3.7 | ISP2 | | White | Rough | Punctiform | Irregular | No | NA | Positive | | Bacteria |
| | PC.L3.9 | DIFCO | | NA | NA | NA | NA | No | NA | Positive | | Bacteria |
| | PC.L3.10 | DIFCO | | NA | NA | NA | NA | No | NA | Positive | | Bacteria |
| | PC.A1.4 | ISP2 | | Yellow | Smooth | Irregular | Irregular | No | NA | Positive | | Bacteria |
| *Peltigera* | PC.N4.4 | DIFCO | 72h | White | Rough | Irregular | Irregular | Yes | *S. aureus* | Positive | | Bacteria |
| | PC.N4.6 | DIFCO | | Cream | Rough | Irregular | Irregular | No | NA | Positive | | Bacteria |
| | PC.N4.7 | ISP2 | 48h | Cream | Rough | Punctiform | Irregular | No | NA | Positive | | Bacteria |
| | PC.N4.8 | DIFCO | 72h | Cream | Rough | Punctiform | Irregular | Yes | *A.baumannii* | Positive | | Bacteria |
| | PC.N4.9 | ISP2 | 4 Days | White | Rough | Irregular | Irregular | No | NA | Positive | | Bacteria |
| | PC.C3.17 | GAUSE | | Yellow | Rough | Punctiform | Irregular | No | NA | Positive | | Bacteria |
| | PC.C3.18 | ISP2 | 72h | Brown | Rough | Filamentous | Irregular | Yes | *S. aureus, C. albicans* | Negative | Positive | Fungi |
| | PC.C3.19 | GAUSE-O | | White | Rough | Circular | Irregular | No | NA | Positive | | Bacteria |
| | PC.C3.20 | DIFCO | | Pink | Rough | Irregular | Irregular | Yes | *S. aureus, A. baumannii* | Positive | | Bacteria |
| | PC.C3.21 | DIFCO | | Yellow | Rough | Punctiform | Irregular | Yes | *S. aureus, A. baumannii, C. albicans* | Positive | | Bacteria |
| | PC.C3.24 | Difco (G) | | White | Rough | Punctiform | Irregular | No | NA | Positive | | Bacteria |
| | PC.C3.25 | GAUSE | | NA | NA | NA | NA | No | NA | Positive | | Bacteria |

*Due to small amount of lichen thalli, the sample genera could not be throughly corroborate

DIFCO: Actinomycete Isolation Agar
GAUSE: Gause Synthetic Agar
GAUSE-O: Gause Oligotrophic Agar
ISP2: Streptomyces Project Medium-2

Description code name: PC corresponds to Chingaza Park; The following letter represents the sublocation point with the number of lichen thalli; Following number represents the numeration for the isolate
