## Supplemental Figures for "The microbiomes of seven lichen genera reveal host specificity, a reduced core community and potential as source of antimicrobials"

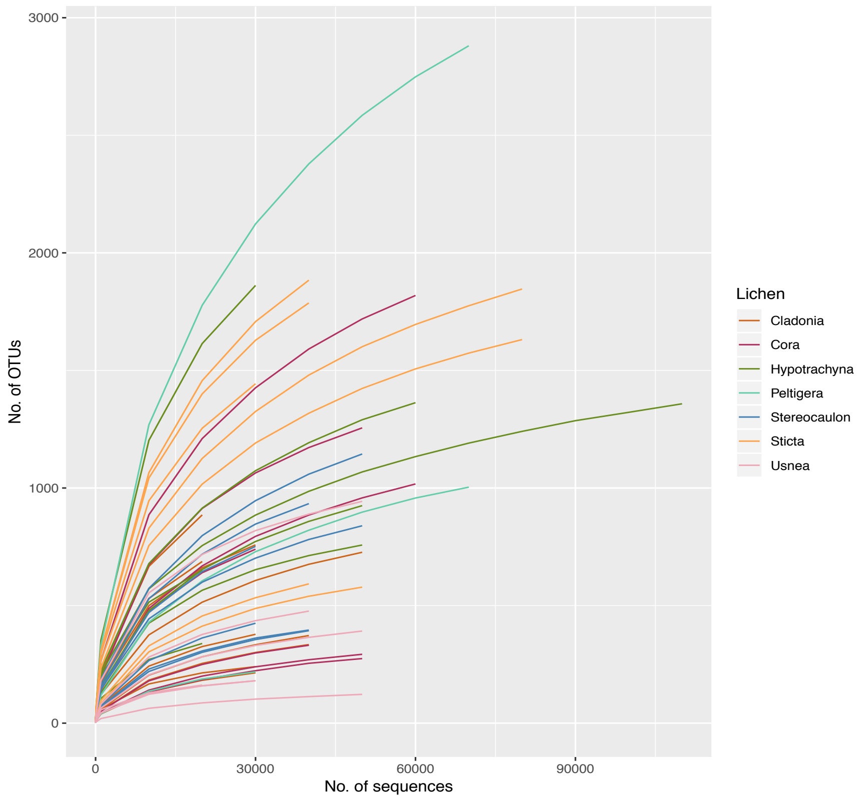


**Figure S1.** Richness of individual samples from microbial communities in lichens. Rarefaction curves of 16S rRNA gene diversity are colored by genera of Lichens.

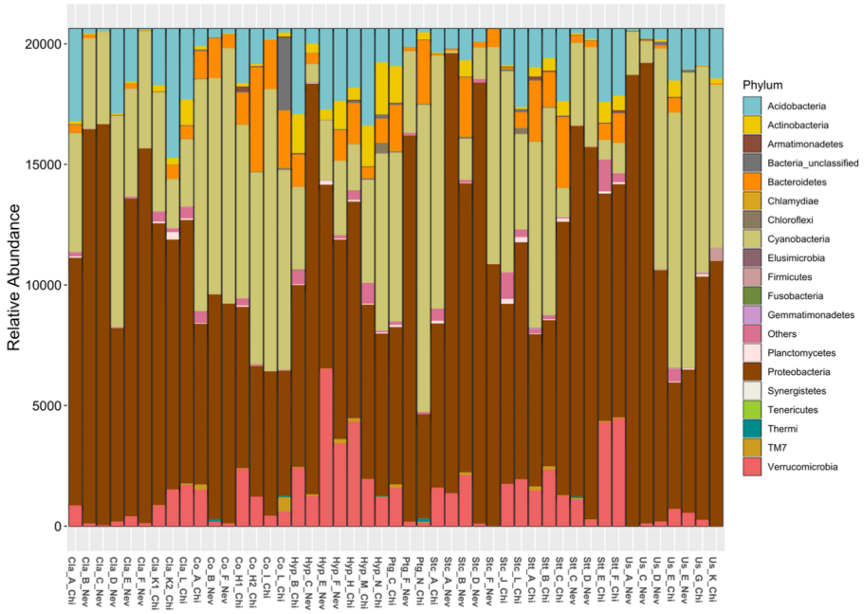


**Figure S2.** Relative abundance at Phylum level of all lichen samples corresponding to *Cladonia* (Cla), *Cora* (Co), *Hypotrachyna* (Hyp), *Peltigera* (Ptg), *Stereocaulon* (Stc), *Sticta* (Stt), *Usnea* (Us). Localities are marked as Nevados (Nev) and Chingaza (Chi).

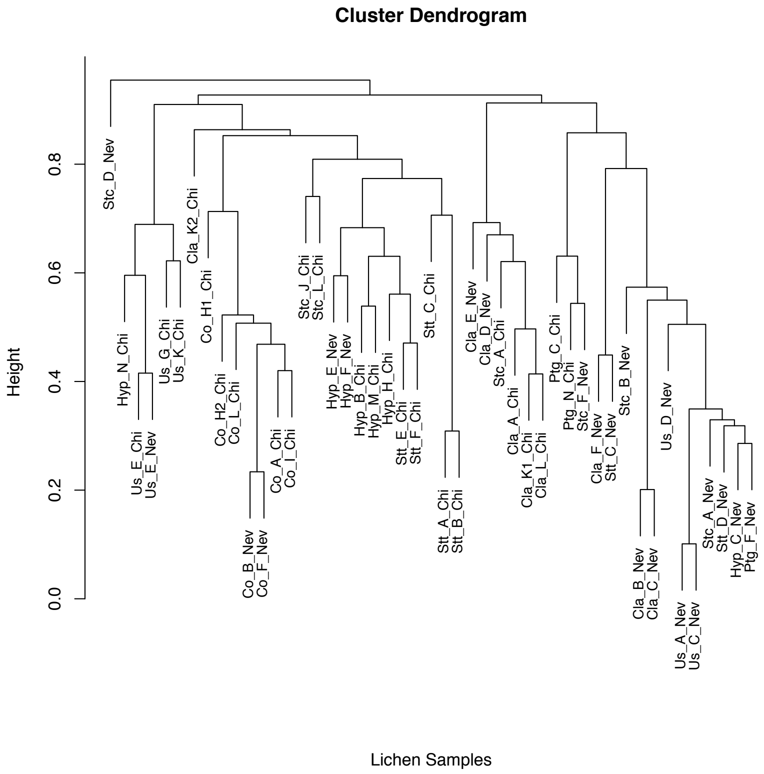


**Figure S3.** Cluster dendrogram of similarity between lichen samples corresponding to *Cladonia* (Cla), *Cora* (Co), *Hypotrachyna* (Hyp), *Peltigera* (Ptg), *Stereocaulon* (Stc), *Sticta* (Stt), *Usnea* (Us) from Nevados (Nev) and Chingaza (Chi) locations. Dendrogram was generated from a Bray-Curtis distance matrix of all samples and the WPGMA hierarchical clustering method.

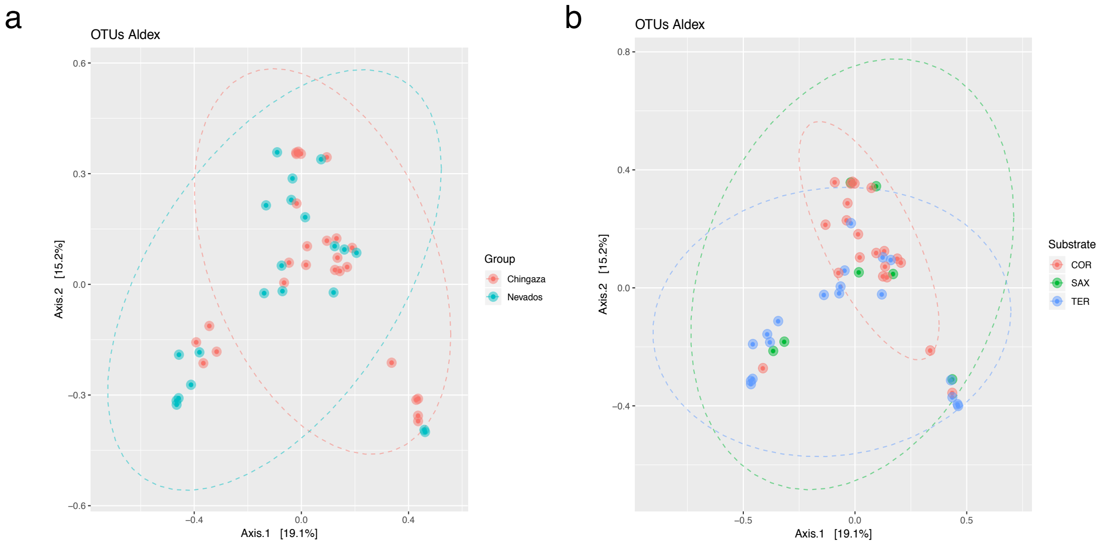


**Figure S4**. Principal coordinates analysis (PCoA) of lichen samples grouped by (a) Localities Chingaza and Nevados; (b) Substrate corresponding to Corticolous (COR), Saxicolous (SAX), Terricolous (TER).

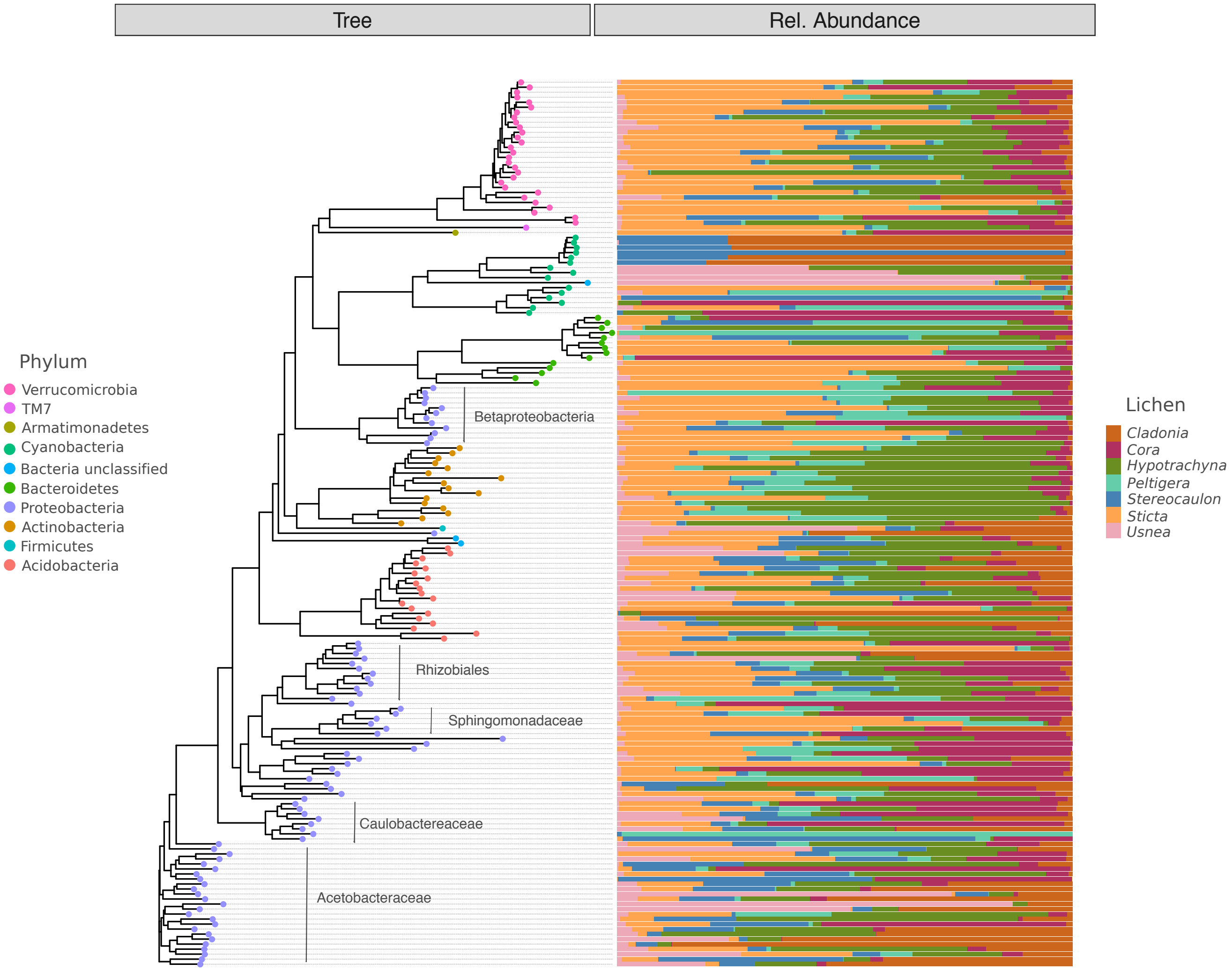


**Figure S5.** Neighbor-Joining tree of 177 significantly differentially abundant OTUs as determined by ALDEx2 and their relative abundance in the seven Lichen genera. The taxonomic clusters of Proteobacteria taxa are indicated.

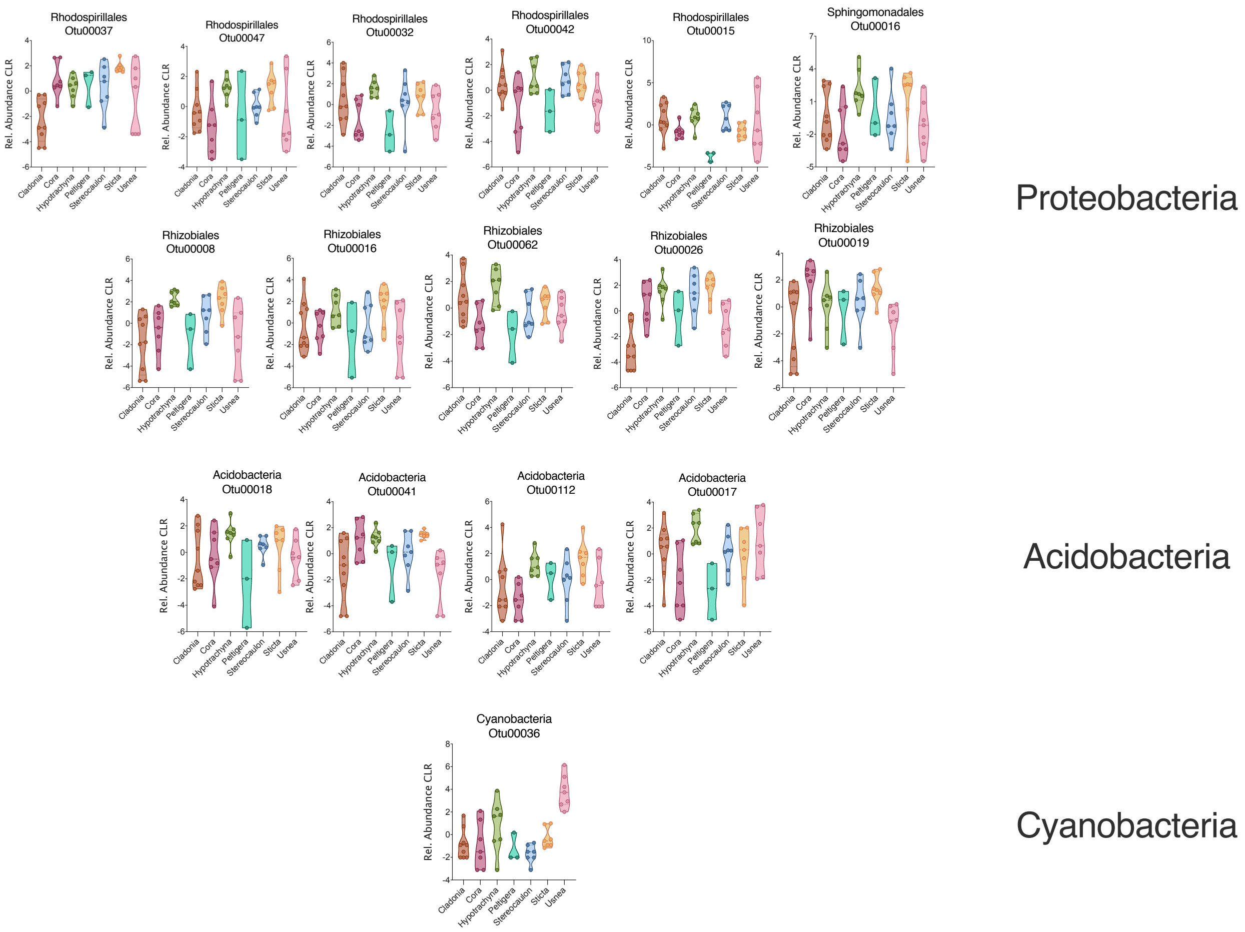


**Figure S6.** Relative abundance (center log transformed, CLR) of the 16 taxa representing the lichen core microbiome.

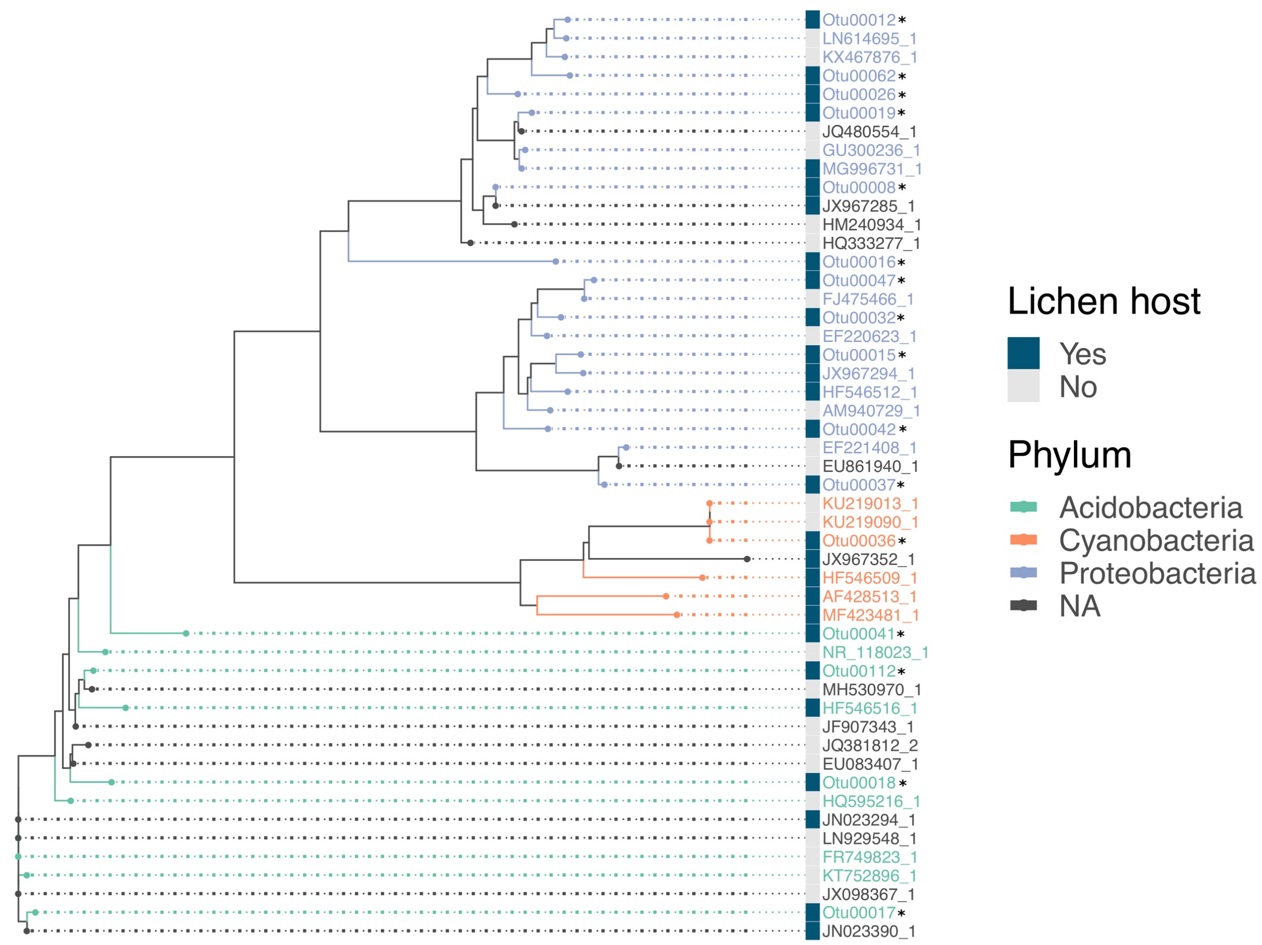


**Figure S7.** Neighbor-joining tree of lichen core OTUs and reference sequences with identity values >98% from GenBank. Bacterial taxa isolated from lichen hosts are marked in blue. Our core OTUs are marked with an asterisk. Each sequence is colored by phylum, sequences that are unclassified are mark as NA.

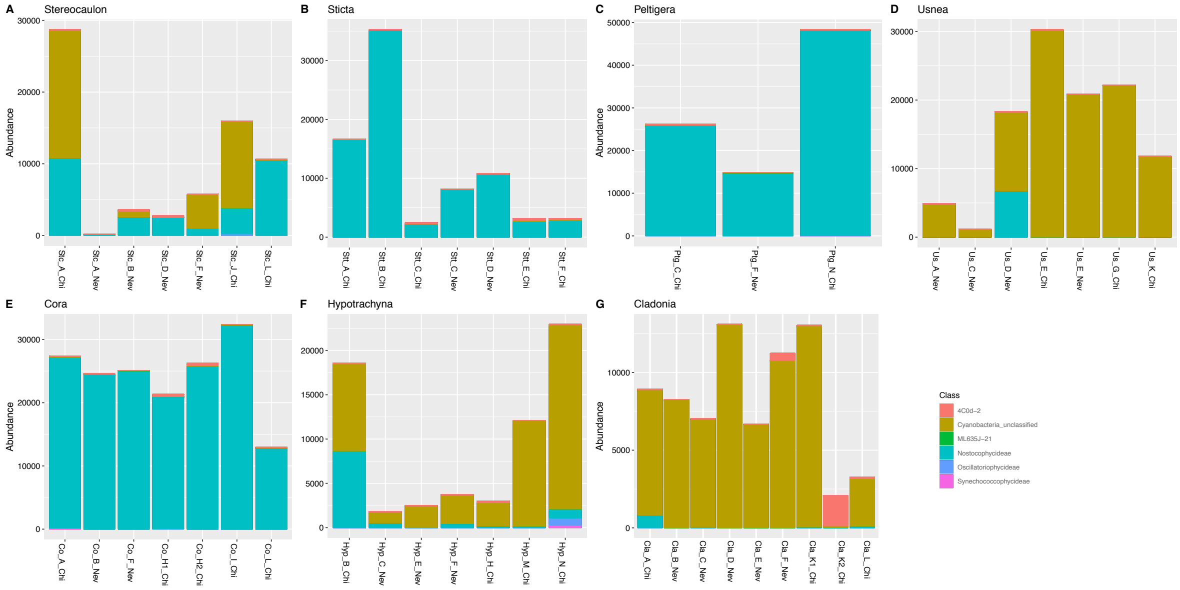


**Figure S8.** Abundance of Cyanobacteria at Class level of lichen samples corresponding to (A) *Stereocaulon* (Stc), (B) *Sticta* (Stt), (C) *Peltigera* (Ptg), (D) *Usnea* (Us), (E) *Cora* (Co), (F) *Hypotrachyna* (Hyp), (G) *Cladonia* (Cla). Localities of sampling are marked as Nevados (Nev) and Chingaza (Chi).
